# From Growth Inhibition to Fluctuations: Testing an Ising Model of Bacterial Responses to Antibiotics

**DOI:** 10.64898/2026.11.23.768705

**Authors:** Pietro Cicuta, Morten Kals, Nir S. Gov

**Affiliations:** Cavendish Laboratory, University of Cambridge, CB3 0HE Cambridge, United Kingdom; Department of Chemical and Biological Physics, Weizmann Institute of Science, Rehovot 7610001, Israel; Department of Physiology, Development and Neuroscience, University of Cambridge, Cambridge CB2 3DY, United Kingdom

## Abstract

Complex networks of biochemical reactions have been proposed to exhibit emergent cooperativity that can be described by an effective Ising model, with a small number of parameters encoding collective interactions. Here this mapping is experimentally tested using the response of three bacterial species to a range of antibiotics acting on different cellular targets. The Ising description is compared to the conventional Hill model, examining both the mean growth-rate response and, importantly, its population growth-rate heterogeneity (PGRH). While both models describe the mean growth-rate dependence on antibiotic concentration comparably well, we find that the Ising model provides a markedly better description of PGRH, offering an additional and more stringent test of the collective description. The fitted Ising interaction strength is systematically related to the Hill exponent and increases with the functional distance of the antibiotic target from the protein-production machinery. These results provide experimental support for describing complex biochemical reaction networks through effective cooperative Ising interactions, and suggest that the collective response of bacterial growth to antibiotics reflects the organization of the underlying metabolic network.

---

There is a growing body of work studying the over-all effective cooperativity in networks of chemical reactions, and how the structure of the network can come into play together with the properties of the nodes [1–6]. This emergent cooperativity was mapped onto a mean-field Ising model [5], in which a single effective coupling parameter determines the Hill coefficient, covering from non-cooperative Michaelis–Menten kinetics up to a switch-like critical point. In the present work we explore and test this mapping using the measured growth-rate response, and its variability, of different species of bacteria to a variety of antibiotics [7].

The effect of different antibiotics, as function of concentration, on the growth rate of several bacteria was recently explored in several recent studies [7–9]. In all these studies it is found that the growth rate decreases, as function of increasing antibiotic concentration, as expected. The growth-rate is observed to undergo a sharp drop at some critical antibiotic concentration, with the sharpness of the drop usually interpreted as an indication for the cooperativity between the metabolic nodes of the bacterial biochemical network that have been perturbed. The data across many antibiotics, with different molecular mechanisms, shows shallow responses for protein-synthesis inhibitors, and increasingly steep responses for antibiotics that act on mechanisms more remote from protein synthesis, up to membrane-synthesis inhibitors, for which the response is close to switch-like. Merchan and Nemenman [6] argue that densely interacting networks with many-body interactions can often be described well by a pairwise maximum-entropy (Ising) model. In a ‘systems’ view of the bacterial cell the observed trend of increasing steepness of the antibiotic response curve with increasing functional distance from the protein production hub could then be interpreted as resulting from the drug perturbation cascading though increasing number of intervening levels, analogous to the signaling cascades described by Ferrell [1].

The step-like behavior of the growth-rate as function of the antibiotic concentration is often phenomenologically fitted to a Hill function, with differing values of the Hill exponent [7]. The Hill function is usually interpreted as an expression for the average occupation of some binding site, where the exponent *n*_*H*_ represents the cooperativity of binding. Hence, for this situation it could be interpreted as the antibiotic binding to its target, which then diminishes the growth rate. While the Hill function description is simple and can be fitted to the data [7], it does not easily correspond to a mechanistic description of the process [10]. Large Hill coefficients produce ultrasensitive, switch-like responses that often require hidden mechanisms such as allosteric coupling, multistep regulation, feedback, or energy-consuming nonequilibrium processes [11, 12], rather than passive thermodynamic equilibrium. In addition, the Hill coefficient is generally only an effective steepness parameter that does not directly correspond to a specific number of binding sites or molecules.

In addition to the growth rate, the population growth rate heterogeneity (PGRH) was measured in [7], where PGRH is defined as the standard deviation of growth rates across many small clonal colonies. There is currently no established theoretical description of the growth rate heterogeneity (PGRH), as function of the antibiotic concentration.

In this work we consider this large dataset of bacterial responses, in terms of both Ising cooperativity or Hill descriptions. We show that (1) both descriptions capture the inflection region of the growth-rate very well; (2) the descriptions are distinguishable, and the analytical link between the Hill and Ising parameters holds as a lower bound in the experimental data; (3) the variability in the growth-rate arises naturally in the Ising framework, and the PGRH data is described better by the Ising model than it is by assuming concentration fluctuations in the background of a Hill description.

## Growth-rate heterogeneity due to concentration fluctuations

One natural source of heterogeneity can arise from fluctuations in the concentration of antibiotics inside the cells. The concentration fluctuations can arise from fluctuations in the number of channels through which the antibiotics enter the cell, the activity of pumps that transports the antibiotics out of the cell, as well as processes that sequester the antibiotic molecules (removing them from the cytoplasm).

Antibiotic concentration fluctuations result in the following estimate for the variance in the growth rate

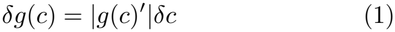

where *g*(*c*) is the average growth-rate at the antibiotic concentration *c, g*(*c*)^*′*^ = *dg*(*c*)*/dc* and *δc* is the variance in the concentration. Concentration fluctuations usually follow Poissonian statistics, such that 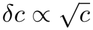. We therefore expect the PGRH that is driven by concentration fluctuations to behave as

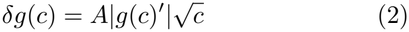

Where *A* is an arbitrary multiplication factor.

The growth-rate is described here by a modified Hill function, where we included an effective “background concentration”, such that: *c → c* + *c*_0_

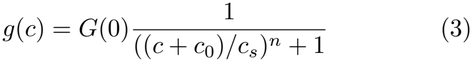

This shift describes a small background concentration of inhibitory effectors that act to limit the growth-rate even in the absence of added antibiotics. This allows the calculated PGRH to have a slower decay in the limit of zero antibiotic concentration, and gives a better fit to the experimental data.

### Ising model for the growth-rate and PGRH

Motivated by the shortcomings of the phenomenological description of the growth-rate dependence on the antibiotic concentration in terms of Hill function, we explore an alternative description of the growth rate and its variance. We consider the biochemical network to be described by a finite number *N* of interacting nodes. The complex cascade of interaction is mapped to an effective interaction between nodes, described by a pair-wise Ising form [13, 14]

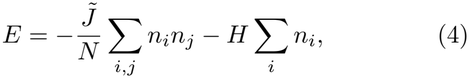

where each of the nodes represents a specific molecular module, such as ion channels or ribosomes, and is modelled to switch between two activity states, *n*_*i*_ (for the *i*-th agent), *n*_*i*_ = 0, 1 denoting the inactive/active state respectively. The first term in Eq.4 describes the effective interactions as an all-to-all spin-spin coupling 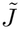. From now we define a reduced interaction parameter to be 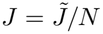. This interaction between the active units can be mediated by complex biochemical and biomechanical circuits within the cell, through competition over limited resources or by direct protein-protein interactions (as in clusters of ion channels on the membrane). For *J >* 0, there is positive feedback between the agents so that as the order parameter *m* (the probability of an agent to be active) increases, the transition rate to state 1 increases (Eq.S5). For *J <* 0 the agents exert a negative feedback on each other.

The second term in Eq.4 denotes the effect of the net metabolic activation field *H* (all energies scaled by some effective thermal energy). This field describes the activating effects of metabolites (such as ATP) and other substrates that maintain activity and drive a positive *H*, which drives the agents to be in the ON (1) state. The binding of antibiotics decreases this field and acts to deactivate the agent, according to Eq.S3, so we write

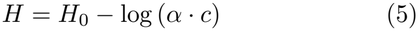

where *c* is the concentration of antibiotics, *α* is some constant that depends on the affinity of the antibiotics and the agent, as well as the antibiotic’s potency. The activation field has a large value *H*_0_ in the absence of antibiotics, such that it gives a large proportion of active agents in the absence of antibiotics.

The steady-state solution for the average probability of an agent to be active, as in the mean-field theory of ferromagnetism, is given by

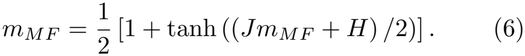

When *J* = 0, (6) yields the sigmoidal curve *m*_*MF*_ = (1*/*2)(1 + tanh(*H/*2)). Increasing positive *J* produces a steeper sigmoid, until a critical value of *J*_*c*_ = 4, where a discontinuity in *m*_*MF*_ appears around *m*_*MF*_ = 1*/*2.

We relate the the proportion of active cellular agents *m*_*MF*_ to the observed growth-rate. For this comparison we have two model parameters to fit, namely *α* (Eq.5) and *J* (Eq.4). The parameter *α* simply shifts the sigmoidal solution *m*_*MF*_ along the concentration axis to fit to the concentration where the growth rate has the maximal gradient, and *J* is varied to fit to the maximal steepness of the growth rate.

We next calculate the variance in the activity *m*_*MF*_ from our spin model [13, 14]

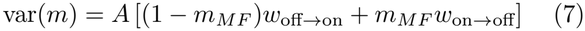

Note that this variance (Eq.7) is distinct from that which arises due to concentration fluctuations (Eq.2). It arises from the dynamics of the cellular agents (spins) turning on/off which gives variations in the instantaneous number of active elements, and is therefore inversely proportional to the total number of these agents: *A ∝* 1*/N* [13].

### Comparison to experimental results

We fitted the experimental data of the growth-rate to both the Hill model (Eq.3) and the Ising model (Eq.6), where we identify the average magnetization *m* as representative of the growth rate. The fitting is done by scanning over parameter space and minimizing the global error function, as described in the SI. Note that both the Hill and Ising models have three free parameters when fitting the growth-rate data, and a single multiplication factor for the fit to the PGRH.

Following the fitting of the growth-rate, we fit the corresponding (species and antibiotic) PGRH data with only one free parameter, the amplitude. For the Hill function this multiplicative constant *A* appears in Eq.2, and for the Ising model in Eq.7.

In Fig.1 we show several typical comparisons between the Hill and Ising models for four antibiotics acting on *E*.*coli* bacteria. These four cases span the range of fitted values of the Hill exponent and Ising interaction. The rest of the comparisons are given in Figs.S4,S5 (all the fitting parameters are given in Tables S2,S3).

**FIG 1.**
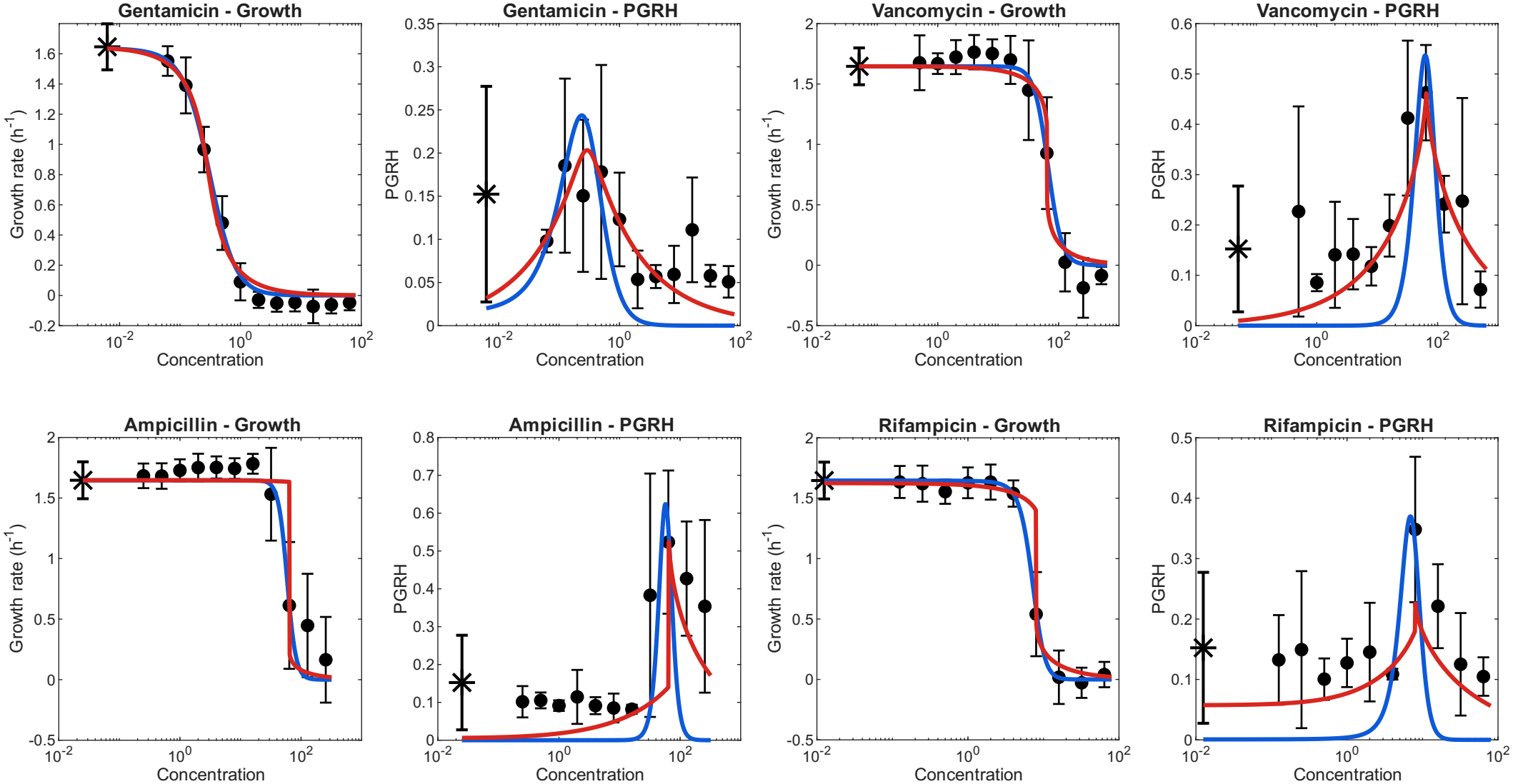
Comparison of Hill (blue) and Ising model (red) fits for Gentamicin, Vancomycin, Ampicillin, and Rifampicin, for *E. coli*. For each antibiotic, the left panel shows the mean growth rate and the right panel shows PGRH. Blue curves represent the Hill model, while red curves represent the Ising model. Experimental measurements are shown in black. The star symbol denote the value at zero antibiotic concentration, plotted at a tenth of the minimal concentration data point in each case. The fits to the other antibiotics are shown in Figs. S4, S5.

The relationship between the fitted Hill exponent and Ising interaction strength, using all three bacteria species, is shown in Fig. 2 (the graphs for each species separately are given in Fig.S10). The solid line is a polynomial fit for the overall correlation, using the relation: *J* = *a·*(*n*_*H*_ − *n*_0_)^*b*^. We find that the limit of non-interacting Ising spins (*J →* 0) corresponds to independent (linear) Hill binding kinetics (*n*_*H*_*→* 1). As the Ising interaction approaches the critical value of *J → J*_*c*_ = 4, the Ising model predicts a sharp drop in the growth-rate, and indeed around this value the corresponding Hill exponents increase sharply. The correspondence between these two descriptions was previously discussed [4], and one can obtain a simple analytic relation if the slope of both the Hill (Eq.3) and Ising (Eq.6) models are equated at their inflection point, giving:

**FIG 2.**
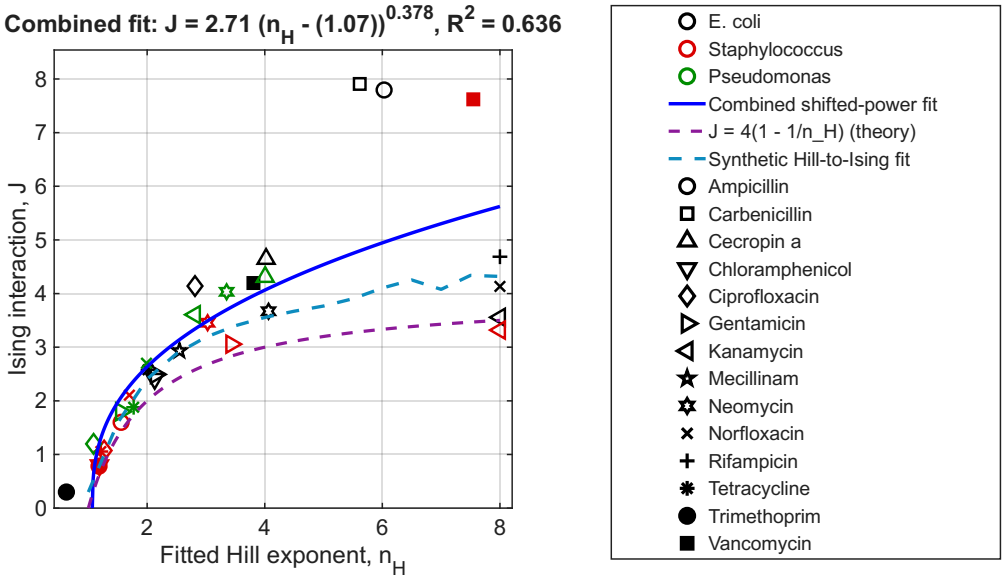
Comparison of fitted Hill exponents (*n*_*H*_) and Ising interaction strengths (*J*) for *E. coli* (black), *Staph* (red) and *Pseu* (green). The different symbols denote the different antibiotics, as given by the legend. The solid line gives a polynomial fit to the data. The purple dashed line gives the prediction of the analytic relation that matches the slope of the Hill function and Ising MF solution at the inflection point [5] (Eq.8). The blue dashed line gives the values of fitting the Ising model to exact Hill functions.

**FIG 3.**
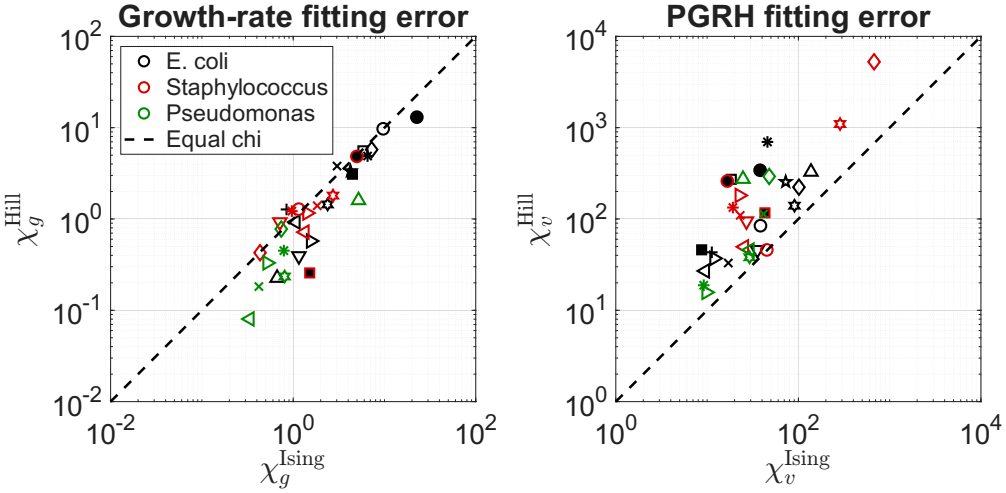
Comparison of fitting errors for the Hill and Ising models for the measured growth-rate (left) and the PGRH (right), for *E. coli* (black), *Staph* (red) and *Pseu* (green). The different symbols denote the different antibiotics, as given by the legend (data symbols as in Fig.2).

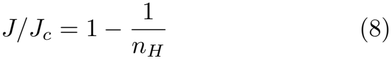

We find that this relation behaves as a lower bound on the results we obtain from fitting the experimental data. This may indicate that to obtain a better fit to the data away from the inflection point, the Ising interaction needs to be higher than given by Eq.8. This is indeed shown by fitting the Ising model to exact Hill functions (see SI section S9), which gives the points along the blue dashed line in Fig. 2, which correspond very well to the experimental data fitting. A few outliers with very high fitted values of *J* are caused by the lack of sufficiently dense experimental data near the inflection point.

Note that there are no cases where the best fit is for *J <* 0, which means that all the cellular targets of the antibiotics effectively have ferromagnetic-like (mutually enhancing) interactions. Second, there is a positive correlation, whereby larger Hill exponents correspond to larger values of the Ising interaction. This means that the interpretation of large Hill exponent as an indication of cooperative interactions, agrees with the Ising model which explicitly describes them.

Ising interactions that are larger than the critical value correspond to the abrupt drop in the growth-rate and the sharp cusp in the PGRH, as shown for example in Fig.1 for Ampicillin and Rifampicin. The abrupt drop occurs since the Ising model enters into a region of bistability, with two coexisting solutions. We show in the SI section S10 that the lower branch solution, which is the one shown in Fig.1 gives a significantly between fit to the PGRH data compared with choosing the higher branch solution (appropriately shifted along the antibiotic concentration axis to fit the sharp drop in the growth-rate, Fig.S12). We do not know if the sharp drops in the growth-rate predicted by the Ising model do actually occur in the cells, or arise here due to fitting an under-sampled experimental data set. However, there is an interesting outcome of this bistability in the Ising model, namely that the PGRH is predicted to be asymmetric around its peak, which seems to appear also in the experimental data, such as for Ampicillin (Fig.1).

In Fig.3 we plot the calculated errors for the fits using the Hill and Ising models (*χ*_*g*_, see SI for details), for all three bacteria and all antibiotics (the graphs for each bacteria separately are given in Fig.S13). For the growth-rate the two models give similar errors, with most of the points along the diagonal. However, for the PGRH, the Ising model gives consistently and significantly better fit of the data (Fig.3), i.e. the shape of the ‘susceptibility’ peak is captured better. These peaks, see Fig.2, have quite different shapes and are systematically shifted in the peak position. For completeness we also tested the fit to the measured PGRH if the concentration fluctuations depend linearly on the concentration, i.e. in Eq.2 we have:*δg*(*c*) = *A*| *g*(*c*)^*′*^| *c*. This did not give significant change to the quality of the fit (Fig.S14), except for one antibiotic.

In [7] it was observed that the Hill exponent grows as the antibiotic acts on a cellular target that is further from the protein synthesis hub. We propose that this can now be interpreted with more insight into the complex networks in the cell, using the Ising model: cellular agents that are more upstream with respect to the protein production, are connected by more biochemical pathways before they affect the protein synthesis. This increase in the number of connecting biochemical pathways results in stronger effective interaction between these elements, corresponding to a larger value of the overall *J* (Eq.4), which reflects a cascade of underlying transitions.

A result of this study is that the effective Ising description accounts not only for the mean growth-rate response but also for the observed population growth-rate heterogeneity. While both the Hill and Ising models reproduce the average growth-inhibition curves, the measured PGRH provides a more discriminating test. In the Hill framework, heterogeneity is estimated from fluctuations in intracellular antibiotic concentration, whereas in the Ising model it emerges naturally from stochastic transitions between the activity states of interacting cellular elements. The Ising description generally provides a substantially better fit to the observed PGRH, particularly near the sharp growth-rate transition. This agreement supports the relevance of collective cellular dynamics to growth-rate fluctuations, although additional sources of heterogeneity, including intracellular antibiotic-concentration fluctuations, may also contribute. This work supports the view that collective properties of the biochemical network shape the cellular response. More broadly, it provides a framework for connecting experimentally accessible response curves and fluctuations to the effective interactions emerging from complex intracellular networks. Future work may establish how these interactions arise from specific biochemical pathways and incorporate them into more detailed models of bacterial growth [15, 16].

## Supporting information

Extra technical details and figures.

