## Supplementary material for "From Growth Inhibition to Fluctuations: Testing an Ising Model of Bacterial Responses to Antibiotics": Extra technical details and figures.

(Dated: November 23, 2026)

### S1. MAPPING LINEAR KINETICS OF ANTIBIOTICS BINDING TO AN ENERGY TERM IN THE ISING MODEL

We start with the linear kinetics of the binding process of the antibiotic to the cellular agent. Let $m(t)$ be the probability of a cellular agent to be in the ON state, equivalent to the magnetization in spin terms. Then

$$\frac{dm}{dt} = -w_{\mathrm{on}\to\mathrm{off}}\, m + w_{\mathrm{off}\to\mathrm{on}}\, (1-m).$$

At steady state, $dm/dt = 0$, giving

$$m^{\mathrm{eq}} = \frac{w_{\mathrm{off}\to\mathrm{on}}}{w_{\mathrm{on}\to\mathrm{off}} + w_{\mathrm{off}\to\mathrm{on}}}.$$

The rates can be described by Glauber dynamics with attempt rate $\gamma$

$$w_{\mathrm{on}\to\mathrm{off}} = \frac{\gamma}{2}\left(1+\tanh\left(\frac{\beta\Delta E}{2}\right)\right), \tag{S1}$$

$$w_{\mathrm{off}\to\mathrm{on}} = \frac{\gamma}{2}\left(1-\tanh\left(\frac{\beta\Delta E}{2}\right)\right). \tag{S2}$$

Using an effective energy field that represents the effect of the antibiotic concentration ($c$)

$$\Delta E \equiv E_{\mathrm{on}} - E_{\mathrm{off}} = k_B T \ln\left(\frac{K_d}{[c]}\right). \tag{S3}$$

and we obtain

$$m^{\mathrm{eq}} = \frac{K_d}{K_d + [c]} = \frac{1}{1+[c]/K_d}.$$

$$p_{\mathrm{off}}^{\mathrm{eq}} = 1 - m^{\mathrm{eq}} = \frac{[c]}{K_d + [c]}.$$

ensuring thermodynamic consistency.

The transition rates between the 0 and 1 states (using Eq.4), are now given as

$$w_{\mathrm{on}\to\mathrm{off}} = \frac{\gamma}{2}\left(1+\tanh\left(\frac{Jm+H}{2}\right)\right), \tag{S4}$$

$$w_{\mathrm{off}\to\mathrm{on}} = \frac{\gamma}{2}\left(1-\tanh\left(\frac{Jm+H}{2}\right)\right). \tag{S5}$$

Here, $J$ and $H$ are dimensionless variables corresponding to $\tilde{J}$ and $\tilde{H}$ scaled by some effective thermal energy.

### S2. CURVE FITTING TO THE HILLS MODEL

#### S2.1. Fitting procedure for the Hill growth model

For each antibiotic, the concentration-dependent mean growth rate and growth-rate heterogeneity can be fitted simultaneously using a Hill-type response model. The model for the mean growth rate was defined as

$$g(c) = \frac{g_0}{1 + \left(\dfrac{c + c_0}{c_{\mathrm{H}}}\right)^{n_{\mathrm{H}}}}, \tag{S6}$$

where $c$ is the antibiotic concentration, $g_0$ is the experimentally measured mean growth rate at zero antibiotic concentration, $c_{\mathrm{H}}$ is the characteristic Hill concentration, $n_{\mathrm{H}}$ is the Hill coefficient, and $c_0$ is a concentration offset.

The three nonlinear parameters fitted to the data were therefore

$$c_{\mathrm{H}}, \qquad n_{\mathrm{H}}, \qquad c_0. \tag{S7}$$

The growth-rate heterogeneity was modeled from the concentration derivative of the Hill curve. The derivative is

$$\frac{dg}{dc} = -\frac{g_0 n_{\mathrm{H}} (c + c_0)^{n_{\mathrm{H}}-1}}{c_{\mathrm{H}}^{n_{\mathrm{H}}} \left[1 + \left(\dfrac{c + c_0}{c_{\mathrm{H}}}\right)^{n_{\mathrm{H}}}\right]^2}. \tag{S8}$$

The predicted heterogeneity was taken to be proportional to the magnitude of this derivative multiplied by the concentration fluctuations which are proportional to $\sqrt{c + c_0}$,

$$v_{\mathrm{model}}(c) = A_{\mathrm{H}} \left|\frac{dg}{dc}\right| \sqrt{c + c_0}, \tag{S9}$$

where $A_{\mathrm{H}}$ is an arbitrary multiplicative scale factor. For each trial set of $(c_{\mathrm{H}}, n_{\mathrm{H}}, c_0)$, the optimal value of $A_{\mathrm{H}}$ was calculated analytically by weighted least squares and was therefore not scanned as an additional nonlinear parameter.

#### S2.2. Objective function

The mean growth-rate and heterogeneity data were fitted simultaneously by minimizing a combined weighted error function.

For the mean growth-rate data, the normalized residual at concentration $c_i$ was defined as

$$R_{g,i} = \frac{g_i - g_{\mathrm{model},i}}{\sigma_{g,i}}, \tag{S10}$$

where $g_i$ is the experimentally measured mean growth rate and $\sigma_{g,i}$ is the experimental uncertainty assigned to the growth-rate measurement.

Each growth-rate residual was additionally weighted by the magnitude of the measured growth rate,

$$w_{g,i} = \frac{|g_i|}{\langle |g| \rangle}. \tag{S11}$$

where

$$\langle |g| \rangle = \frac{1}{N} \sum_{j=1}^{N} |g_j|, \tag{S12}$$

and $N$ is the total number of measured concentration points, including zero concentration. This normalization ensures that the average weight is unity, giving greater importance to measurements with higher growth rates.

The growth-rate contribution to the objective error function was therefore

$$\chi_g = \sum_i w_{g,i} R_{g,i}^2. \tag{S13}$$

For the growth-rate heterogeneity data, the normalized residual was

$$R_{v,i} = \frac{v_i - v_{\mathrm{model},i}}{\sigma_{v,i}}, \tag{S14}$$

where $v_i$ is the measured growth-rate heterogeneity and $\sigma_{v,i}$ is its experimental uncertainty.
The he heterogeneity residuals were weighted by the measured heterogeneity data,

$$w_{v,i} = \frac{|v_i|}{\langle |v| \rangle}. \tag{S15}$$

The heterogeneity contribution was then

$$\chi_v = \sum_i w_{v,i} R_{v,i}^2. \tag{S16}$$

The most general total objective function minimized by the fit can be written as

$$\chi_{\mathrm{tot}} = W_g \chi_g + W_v \chi_v, \tag{S17}$$

However, we used only the growth-rate weighting

$$W_g = 1, \qquad W_v = 0. \tag{S18}$$

Other weighting, for example giving equal weight to both data sets, led to very large errors in fitting the growth-rate data, for which there is much greater confidence. This is shown in Figs.S1,S2.

#### S2.3. Determination of the heterogeneity scale factor

For a given set of Hill parameters, define

$$s_i = \left| \frac{dg}{dc} \right|_{c_i} \sqrt{c_i + c_0}. \tag{S19}$$

The heterogeneity model is then

$$v_{\mathrm{model},i} = A_{\mathrm{H}} s_i. \tag{S20}$$

For each trial set of $(c_{\mathrm{H}}, n_{\mathrm{H}}, c_0)$, the value of $A_{\mathrm{H}}$ minimizing the weighted heterogeneity error was calculated analytically as

$$A_{\mathrm{H}} = \frac{\sum_i \left( \frac{w_{v,i}}{\sigma_{v,i}^2} \right) s_i v_i}{\sum_i \left( \frac{w_{v,i}}{\sigma_{v,i}^2} \right) s_i^2}. \tag{S21}$$

The fitted scale factor was restricted to be non-negative.

#### S2.4. Initial parameter scan

An initial three-dimensional scan was performed over $c_{\mathrm{H}}$, $n_{\mathrm{H}}$, and $c_0$.
Because the concentration ranges differed substantially between antibiotics, the scan limits for $c_{\mathrm{H}}$ and $c_0$ were defined separately for each antibiotic. Let $c_{\mathrm{min}}$ denote the smallest non-zero experimental concentration and $c_{\mathrm{max}}$ the largest experimental concentration.
The Hill concentration was scanned over

$$0.1\, c_{\mathrm{min}} \le c_{\mathrm{H}} \le 2\, c_{\mathrm{max}}. \tag{S22}$$

A total of 55 logarithmically spaced values of $c_\mathrm{H}$ were evaluated.

The Hill coefficient was scanned linearly over

$$0.3 \leq n_\mathrm{H} \leq 8, \tag{S23}$$

using 55 equally spaced values. The corresponding spacing was

$$\Delta n_\mathrm{H} = \frac{8 - 0.3}{55 - 1} \simeq 0.1426. \tag{S24}$$

The concentration offset was scanned over

$$10^{-4} c_\mathrm{min} \leq c_0 \leq c_\mathrm{max}, \tag{S25}$$

using 55 logarithmically spaced values.

The initial coarse scan therefore evaluated

$$55^3 = 166375 \tag{S26}$$

parameter combinations for each antibiotic.

For every parameter combination, the Hill growth curve and its analytical derivative were calculated at all experimental concentrations, the optimal heterogeneity scale factor $A_\mathrm{H}$ was determined analytically, and the combined objective function $\chi_\mathrm{tot}$ was evaluated.

The parameter combination with the smallest value of $\chi_\mathrm{tot}$ was retained as the best result of the coarse scan.

#### S2.5. Continuous parameter refinement

The best point from the coarse parameter scan was subsequently used as the starting point for a continuous local refinement using MATLAB's `fminsearch` routine.

All three positive Hill parameters were optimized in logarithmic coordinates,

$$p_1 = \ln c_\mathrm{H}, \qquad p_2 = \ln n_\mathrm{H}, \qquad p_3 = \ln c_0. \tag{S27}$$

This parametrization guarantees that all fitted parameters remain positive during the optimization.

The continuous refinement was restricted to remain inside the parameter ranges used in the initial scan. Trial values outside these limits were assigned a large penalty in the objective function.

At every step of the refinement, the Hill curve was recalculated, its analytical derivative was evaluated, the optimal heterogeneity scale factor $A_\mathrm{H}$ was recalculated analytically, and the complete weighted objective function was evaluated.

The final reported values of

$$c_\mathrm{H}, \qquad n_\mathrm{H}, \qquad c_0, \qquad A_\mathrm{H} \tag{S28}$$

therefore correspond to the refined minimum of the combined growth-rate and heterogeneity objective function, rather than being restricted to the discrete values used in the initial parameter scan.

### S3. CURVE FITTING TO THE ISING MODEL

#### S3.1. Fitting procedure for the Ising-like growth model

For each antibiotic, the concentration-dependent mean growth rate and growth-rate heterogeneity were fitted simultaneously using an Ising-like model. The model is controlled by three parameters, the interaction strength $J_0$, the concentration-scaling parameter $\alpha$, and the concentration offset $c_0$.

For a given antibiotic concentration $c$, the effective field was defined as

$$h(c) = 4 - \ln\left[(c + c_0)\alpha\right]. \tag{S29}$$

The magnetization $m$ was obtained numerically from the steady-state condition of the Ising-like dynamics. The transition rates were

$$r_+ = \frac{1}{1 + \exp[-(J_0 m + h)]}, \tag{S30}$$

and

$$r_- = \frac{1}{1 + \exp[J_0 m + h]}. \tag{S31}$$

The corresponding dynamical equation is

$$\frac{dm}{dt} = (1 - m)r_+ - mr_-. \tag{S32}$$

At steady state, this reduces to

$$m = r_+. \tag{S33}$$

The steady-state magnetization was therefore calculated iteratively at each drug concentration. The iteration was started from $m = 0$ and continued until the change in magnetization was smaller than $10^{-10}$, with a maximum of 5000 iterations. A damping factor of 0.5 was used to improve numerical stability.

The predicted mean growth rate was taken to be

$$g_{\text{model}}(c) = g_0 m(c), \tag{S34}$$

where $g_0$ is the experimentally measured mean growth rate at zero antibiotic concentration.

The model prediction for the growth-rate heterogeneity was calculated from

$$v_{\text{Ising}}(c) = \sqrt{r_+(1 - m) + r_- m}. \tag{S35}$$

Because the absolute magnitude of this quantity is not fixed by the model, an additional multiplicative scale factor $A_v$ was introduced,

$$v_{\text{model}}(c) = A_v\, v_{\text{Ising}}(c). \tag{S36}$$

For each trial set of $(J_0, \alpha, c_0)$, the optimal value of $A_v$ was calculated analytically by weighted least squares rather than being scanned as an independent nonlinear parameter.

#### S3.2. Objective function

The fit was determined by minimizing a combined error function containing contributions from both the mean growth-rate data and the growth-rate heterogeneity data. For the mean growth rate, the normalized residual was

$$R_{g,i} = \frac{g_i - g_{\text{model},i}}{\sigma_{g,i}}, \tag{S37}$$

where $g_i$ is the experimentally measured mean growth rate and $\sigma_{g,i}$ is the experimental uncertainty assigned to the growth-rate measurement.

Each residual was additionally weighted by the magnitude of the measured growth rate,

$$w_{g,i} = \frac{|g_i|}{\langle |g| \rangle}. \tag{S38}$$

The growth-rate contribution to the fitting error was therefore

$$\chi_g = \sum_i w_{g,i} R_{g,i}^2. \tag{S39}$$

For the growth-rate heterogeneity, the normalized residual was

$$R_{v,i} = \frac{v_i - v_{\text{model},i}}{\sigma_{v,i}}, \tag{S40}$$

where $v_i$ is the measured growth-rate heterogeneity and $\sigma_{v,i}$ is its experimental uncertainty.

The weights were determined from the measured heterogeneity values,

$$w_{v,i} = \frac{|v_i|}{\langle |v| \rangle}. \tag{S41}$$

The heterogeneity contribution to the fitting error was then

$$\chi_v = \sum_i w_{v,i} R_{v,i}^2. \tag{S42}$$

As for the case of fitting the Hill function (Eq.S17) the total objective function minimized by the fit was

$$\chi_{\text{tot}} = W_g \chi_g + W_v \chi_v, \tag{S43}$$

with

$$W_g = 1, \qquad W_v = 0. \tag{S44}$$

#### S3.3. Parameter scan

An initial three-dimensional parameter scan was carried out over $J_0$, $\alpha$, and $c_0$.

The interaction parameter $J_0$ was scanned linearly over the range

$$0.3 \le J_0 \le 8.0. \tag{S45}$$

A total of 49 equally spaced values were used, corresponding to a spacing

$$\Delta J_0 = \frac{8.0 - 0.3}{49 - 1} \simeq 0.1604. \tag{S46}$$

The parameter $\alpha$ was scanned logarithmically using 35 points. Because the experimental concentration range differs between antibiotics, the $\alpha$ range was defined separately for each antibiotic. If $c_{\text{min}}$ and $c_{\text{max}}$ denote the smallest non-zero and largest experimental concentrations, respectively, the scan range was

$$\alpha_{\text{min}} = \frac{0.1}{c_{\text{max}}}, \tag{S47}$$

and

$$\alpha_{\text{max}} = \frac{1000}{c_{\text{min}}}. \tag{S48}$$

Thus, for each antibiotic, 35 logarithmically spaced values of $\alpha$ were evaluated between these limits.

The concentration offset $c_0$ was also scanned logarithmically using 35 points. Its range was defined separately for each antibiotic as

$$c_{0,\text{min}} = 10^{-4} c_{\text{min}}, \tag{S49}$$

and

$$c_{0,\text{max}} = c_{\text{max}}. \tag{S50}$$

Consequently, the initial coarse search evaluated

$$49 \times 35 \times 35 = 60025 \tag{S51}$$

combinations of $(J_0, \alpha, c_0)$ for each antibiotic.

For each parameter combination, the Ising model was solved numerically at all experimentally measured concentrations, the optimal heterogeneity scale factor $A_v$ was determined analytically, and the total objective function $\chi_{\text{tot}}$ was calculated. The parameter combination giving the smallest value of $\chi_{\text{tot}}$ was retained as the best result of the coarse scan.

#### S3.4. Continuous parameter refinement

The best parameter combination obtained from the coarse scan was subsequently used as the starting point for a continuous local optimization using MATLAB's `fminsearch` routine.

During this refinement, $J_0$ was optimized directly, while $\alpha$ and $c_0$ were optimized in logarithmic coordinates,

$$p_1 = J_0, \qquad p_2 = \ln \alpha, \qquad p_3 = \ln c_0. \tag{S52}$$

The logarithmic representation guarantees that $\alpha$ and $c_0$ remain positive.

The refinement was constrained to remain inside the parameter ranges explored by the coarse scan. Trial parameter sets outside these limits were assigned a large penalty in the objective function. At every step of the continuous optimization, the Ising model was recalculated, the optimal heterogeneity scale factor $A_v$ was recalculated analytically, and the complete weighted objective function was reevaluated.

The final reported parameters $(J_0, \alpha, c_0)$ were therefore not restricted to the discrete values of the original scan. The coarse scan was used to locate the appropriate region of parameter space, while the subsequent continuous optimization provided the final refined parameter estimates.

### S4. COMPARISON OF ISING MODEL FITS WITH DIFFERENT ERROR FUNCTION WEIGHTS

To account for local variations in the measured growth rate, the fitting error was weighted using the growth-rate values at each concentration and its two nearest neighboring concentrations. Specifically, for an interior data point $i$, we defined the local average growth rate as

$$\widetilde{g}_i = \frac{|g_{i-1}| + |g_i| + |g_{i+1}|}{3}, \tag{S53}$$

where $g_i$ is the experimentally measured mean growth rate at concentration $c_i$. For the first and last data points, the average was calculated using only the point itself and its single available neighbor.

The normalized weight assigned to each data point was then

$$w_i = \frac{\widetilde{g}_i}{\langle \widetilde{g} \rangle}, \tag{S54}$$

where $\langle \widetilde{g} \rangle$ denotes the average of the local growth-rate values over all measured concentrations.

The weighted fitting error for the mean growth rate was calculated as

$$\chi_g = \sum_i w_i \left( \frac{g_i - g_{\rm model}(c_i)}{\sigma_{g,i}} \right)^2, \tag{S55}$$

where $\sigma_{g,i}$ is the experimental uncertainty in the growth-rate measurement. Importantly, the local averaging was applied only to the weights, while the fitting residuals were calculated using the original, unsmoothed experimental measurements.

While this procedure reduces the sensitivity of the weighting to fluctuations at individual data points, it was most important when fitting the Ising model to a few data sets due to this method giving greater importance to measurements in concentration regions characterized by higher growth rates. As can be seen in Fig.S3a,b this improved the fitting by shifting the sharp jump in the Ising model graph to lower antibiotic concentrations. For all other cases, both methods give very similar fitting parameters, as shown in Table S1.

### S5. COMPARISON OF FITS OF THE HILLS AND ISING FUNCTIONS TO THE EXPERIMENTAL DATA OF *E. COLI.*

In Figs.S4,S5 we give the comparisons of the Hill and Ising models to the experimental data, for the different antibiotics not shown in Fig.1. The fitting parameters for all the antibiotics are given in Table S2 and Table S3, for the Hill and Ising models respectively.

TABLE S1: Comparison of the fitted Ising-model parameters obtained using the regular growth-rate weighting and the three-point neighbor-weighted fitting procedure (Fig.S3). $J_0$ is the interaction strength, $\alpha$ is the concentration-scaling parameter, $c_0$ is the concentration offset, and $A_v$ is the multiplicative scaling factor for the predicted growth-rate heterogeneity. In both procedures, only the growth-rate fitting error was minimized ($W_g = 1$, $W_v = 0$).

| **Antibiotic** | **Fitting method** | $\mathbf{J_0}$ | $\boldsymbol{\alpha}$ | $\mathbf{c_0}$ | $\mathbf{A_v}$ |
|---|---|---|---|---|---|
| Ampicillin | Regular | 5.000 | 4.448 | $4.137 \times 10^{-5}$ | 0.8392 |
| Ampicillin | Neighbor-weighted | 7.795 | 15.538 | 0.1065 | 1.1151 |
| Carbenicillin | Regular | 5.000 | 4.457 | $2.500 \times 10^{-5}$ | 1.0728 |
| Carbenicillin | Neighbor-weighted | 7.908 | 15.820 | 0.03422 | 2.2631 |
| Vancomycin | Regular | 4.090 | 6.564 | $5.000 \times 10^{-5}$ | 0.6924 |
| Vancomycin | Neighbor-weighted | 4.196 | 6.855 | $1.651 \times 10^{-4}$ | 0.7005 |

TABLE S2: Fitted parameters of the Hill model for the different antibiotics. Here $c_H$ is the characteristic Hill concentration, $n_H$ is the Hill exponent, $c_0$ is the concentration offset, and $A_h$ is the multiplicative factor used to fit the growth-rate heterogeneity (PGRH). The fitting was performed using $W_g = 1$ and $W_v = 0$.

| **Antibiotic** | $c_H$ | $n_H$ | $c_0$ | $A_h$ |
|---|---|---|---|---|
| Ampicillin | 58.9365 | 6.0298 | $2.5 \times 10^{-5}$ | 1.91654 |
| Carbenicillin | 42.7270 | 5.61533 | $2.5 \times 10^{-5}$ | 3.87670 |
| Cecropin A | 1.79743 | 4.02253 | 0.157656 | 0.407625 |
| Chloramphenicol | 1.94266 | 2.12842 | 0.131298 | 0.246893 |
| Ciprofloxacin | 0.0165462 | 2.81359 | $1.95 \times 10^{-7}$ | 0.0354153 |
| Gentamicin | 0.329107 | 2.14751 | 0.0262875 | 0.149701 |
| Kanamycin | 5.10063 | 8.00000 | 3.38286 | 0.172658 |
| Mecillinam | 0.826947 | 2.54980 | $1.95 \times 10^{-7}$ | 0.386467 |
| Neomycin | 0.762646 | 4.06429 | 0.230378 | 0.0893916 |
| Norfloxacin | 1.73493 | 8.00000 | 1.12542 | 0.175571 |
| Rifampicin | 11.1439 | 8.00000 | 4.04953 | 0.373598 |
| Tetracycline | 0.738657 | 2.01476 | $1.56 \times 10^{-6}$ | 0.196839 |
| Trimethoprim | 0.509826 | 0.626311 | $3.125 \times 10^{-6}$ | 0.387788 |
| Vancomycin | 66.8485 | 3.80755 | $5.0 \times 10^{-5}$ | 2.75269 |

### S6. COMPARISON OF FITS OF THE HILLS AND ISING FUNCTIONS TO THE EXPERIMENTAL DATA OF *STAPH.*

In Figs.S4,S5 we give the comparisons of the Hill and Ising models to the experimental data, for the different antibiotics not shown in Fig.1. The fitting parameters for all the antibiotics are given in Table S4 and Table S5, for the Hill and Ising models respectively.

### S7. COMPARISON OF FITS OF THE HILLS AND ISING FUNCTIONS TO THE EXPERIMENTAL DATA OF *PSEUDOMONAS.*

In Figs.S8,S9 we give the comparisons of the Hill and Ising models to the experimental data, for the different antibiotics not shown in Fig.1. The fitting parameters for all the antibiotics are given in Table S6 and Table S7, for the Hill and Ising models respectively.

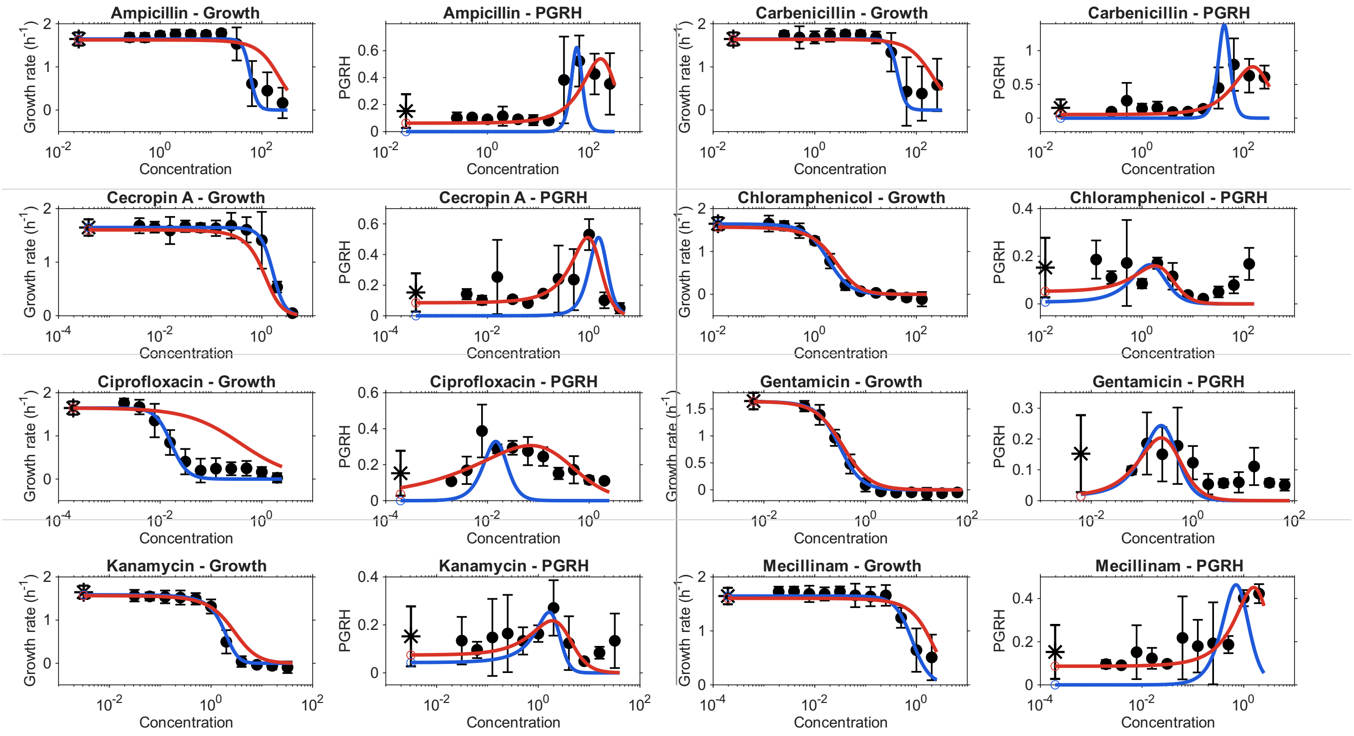


FIG. S1: **Comparison of Hill-model fits to bacterial growth rates and growth-rate heterogeneity (first eight antibiotics).** For each antibiotic, the left panel shows the mean growth rate as a function of antibiotic concentration, while the right panel shows the population growth-rate heterogeneity (PGRH). Black symbols represent the experimental measurements, with error bars indicating the experimental uncertainties. Blue solid curves correspond to fits obtained by minimizing only the growth-rate error ($W_g = 1$, $W_v = 0$), whereas red solid curves correspond to simultaneous fitting of growth rate and PGRH ($W_g = 1$, $W_v = 1$).

TABLE S3: Fitted parameters of the Ising model for the different antibiotics. Here $J_0$ is the effective interaction strength, $\alpha$ is the concentration-scaling parameter, $c_0$ is the concentration offset, and $A_i$ is the multiplicative factor used to fit the growth-rate heterogeneity (PGRH). The growth-rate fitting error was weighted using the local average of three neighboring growth-rate measurements. The fitting was performed using $W_g = 1$ and $W_v = 0$.

| **Antibiotic** | $J_0$ | $\alpha$ | $c_0$ | $A_i$ |
|---|---|---|---|---|
| Ampicillin | 7.79507 | 15.5379 | 0.106475 | 1.11514 |
| Carbenicillin | 7.90770 | 15.8202 | 0.0342227 | 2.26314 |
| Cecropin A | 4.64634 | 256.539 | $3.90625 \times 10^{-7}$ | 0.910229 |
| Chloramphenicol | 2.40967 | 104.224 | $1.25 \times 10^{-5}$ | 0.188290 |
| Ciprofloxacin | 4.14003 | 27440.7 | $2.10563 \times 10^{-6}$ | 0.508554 |
| Gentamicin | 2.49310 | 651.797 | $6.25 \times 10^{-6}$ | 0.287481 |
| Kanamycin | 3.55724 | 164.047 | 0.420246 | 0.253156 |
| Mecillinam | 2.93382 | 266.775 | $1.95 \times 10^{-7}$ | 0.640365 |
| Neomycin | 3.66995 | 660.036 | 0.0104346 | 0.499939 |
| Norfloxacin | 4.13308 | 645.774 | 0.162365 | 0.391894 |
| Rifampicin | 4.68664 | 55.7757 | 1.31499 | 0.357059 |
| Tetracycline | 2.57864 | 262.591 | $1.56 \times 10^{-6}$ | 0.248076 |
| Trimethoprim | 0.30000 | 298.543 | $3.125 \times 10^{-6}$ | 0.412273 |
| Vancomycin | 4.19631 | 6.85493 | $1.65068 \times 10^{-4}$ | 0.700502 |

TABLE S4: Fitted parameters of the Hill model for antibiotics acting on Staphylococcus. Here $c_H$ is the characteristic Hill concentration, $n_H$ is the Hill exponent, $c_0$ is the concentration offset, and $A_h$ is the multiplicative factor used to fit the population growth-rate heterogeneity (PGRH). The fitting was performed with $W_g = 1$ and $W_v = 0$.

| **Antibiotic** | $c_H$ | $n_H$ | $c_0$ | $A_h$ |
|---|---|---|---|---|
| Ampicillin | 0.0433942 | 1.55724 | 0.0026907 | 0.142502 |
| Chloramphenicol | 1.93786 | 1.18903 | 0.0179395 | 0.311659 |
| Ciprofloxacin | 0.111616 | 1.26832 | $1.56 \times 10^{-6}$ | 0.222896 |
| Gentamicin | 0.523863 | 3.43106 | 0.224161 | 0.155704 |
| Kanamycin | 337.842 | 8.00000 | 227.494 | 2.63222 |
| Neomycin | 6.07828 | 3.02801 | $3.125 \times 10^{-6}$ | 0.744778 |
| Norfloxacin | 0.367182 | 1.69730 | $3.125 \times 10^{-6}$ | 0.173554 |
| Tetracycline | 0.0207333 | 1.21225 | $1.31161 \times 10^{-4}$ | 0.0555851 |
| Trimethoprim | 0.786180 | 1.18100 | $3.125 \times 10^{-6}$ | 0.577211 |
| Vancomycin | 0.232749 | 7.54682 | 0.0321128 | 0.109480 |

TABLE S5: Fitted parameters of the Ising model for antibiotics acting on Staphylococcus. Here $J_0$ is the effective interaction strength, $\alpha$ is the concentration-scaling parameter, $c_0$ is the concentration offset, and $A_i$ is the multiplicative factor used to fit PGRH. The growth-rate fitting error was weighted using the local average of three neighboring growth-rate measurements. The fitting was performed with $W_g = 1$ and $W_v = 0$.

| **Antibiotic** | $J_0$ | $\alpha$ | $c_0$ | $A_i$ |
|---|---|---|---|---|
| Ampicillin | 1.59985 | 3024.97 | $6.37185 \times 10^{-4}$ | 0.365473 |
| Chloramphenicol | 0.797820 | 42.2574 | 0.0201506 | 0.106536 |
| Ciprofloxacin | 1.07382 | 842.597 | $1.56 \times 10^{-6}$ | 0.547975 |
| Gentamicin | 3.05664 | 713.552 | 0.0790621 | 0.211718 |
| Kanamycin | 3.31935 | 2.40182 | 22.1676 | 0.285990 |
| Neomycin | 3.45607 | 46.9218 | $3.125 \times 10^{-6}$ | 0.541882 |
| Norfloxacin | 2.10603 | 424.569 | $3.125 \times 10^{-6}$ | 0.247741 |
| Tetracycline | 1.05533 | 4564.11 | $1.60884 \times 10^{-4}$ | 0.163298 |
| Trimethoprim | 0.782754 | 104.075 | $3.125 \times 10^{-6}$ | 0.181277 |
| Vancomycin | 7.62059 | 7758.26 | $3.13025 \times 10^{-6}$ | 0.890342 |

### S8. COMPARISON OF THE ISING INTERACTION PARAMETER $J_0$ AND HILL EXPONENT $n_{hill}$ TO THE EXPERIMENTAL DATA OF THE GROWTH-RATE.

Comparison of fitted Hill exponents and Ising interaction strengths (Fig.S10).

TABLE S6: Fitted parameters of the Hill model for antibiotics acting on Pseudomonas. Here $c_H$ is the characteristic Hill concentration, $n_H$ is the Hill exponent, $c_0$ is the concentration offset, and $A_h$ is the multiplicative factor used to fit the population growth-rate heterogeneity (PGRH). The fitting was performed with $W_g = 1$ and $W_v = 0$.

| **Antibiotic** | $c_H$ | $n_H$ | $c_0$ | $A_h$ |
|---|---|---|---|---|
| Cecropin A | 7.66962 | 4.00720 | $6.25 \times 10^{-6}$ | 0.666977 |
| Ciprofloxacin | 0.0671101 | 1.08862 | $3.90625 \times 10^{-7}$ | 0.274307 |
| Gentamicin | 0.348592 | 1.55534 | 0.0132142 | 0.220277 |
| Kanamycin | 8.34651 | 2.81205 | 0.722343 | 1.52324 |
| Neomycin | 3.07602 | 3.35123 | $1.5625 \times 10^{-6}$ | 0.656247 |
| Norfloxacin | 3.40274 | 1.98838 | $7.8125 \times 10^{-7}$ | 0.941797 |
| Tetracycline | 4.97791 | 1.77014 | 0.499027 | 0.723106 |

TABLE S7: Fitted parameters of the Ising model for antibiotics acting on Pseudomonas. Here $J_0$ is the effective interaction strength, $\alpha$ is the concentration-scaling parameter, $c_0$ is the concentration offset, and $A_i$ is the multiplicative factor used to fit PGRH. The growth-rate fitting error was weighted using the local average of three neighboring growth-rate measurements. The fitting was performed with $W_g = 1$ and $W_v = 0$.

| **Antibiotic** | $J_0$ | $\alpha$ | $c_0$ | $A_i$ |
|---|---|---|---|---|
| Cecropin A | 4.30979 | 56.8104 | 0.0612377 | 0.317610 |
| Ciprofloxacin | 1.19797 | 1739.23 | $6.97631 \times 10^{-4}$ | 0.581538 |
| Gentamicin | 1.79765 | 424.222 | 0.00130184 | 0.270968 |
| Kanamycin | 3.60624 | 42.3346 | $2.5 \times 10^{-5}$ | 0.464707 |
| Neomycin | 4.03252 | 111.301 | $1.5625 \times 10^{-6}$ | 0.331152 |
| Norfloxacin | 2.70176 | 61.7470 | $7.8125 \times 10^{-7}$ | 0.468227 |
| Tetracycline | 1.88093 | 31.7356 | 0.104785 | 0.216291 |

### S9. FITTING THE ISING MODEL TO EXACT HILL FUNCTION

To get a better understanding of the relation between the Hill function exponent and the corresponding Ising model interaction, we produced data sets of exact Hill functions of different Hill exponent, densely and equally spaced along the logarithmic concentration axis. We then used the same procedure described in section S2 to fit it to the Ising model. The results are shown in Fig.S11, and the fitted values of $J_0$ are plotted as the blue dashed line in Fig.2.

We see clearly that as the Ising model becomes sharper to match the Hill function for larger values of the exponent $n_H$, the two functions become more and more different as they approach the inflection point (Fig.S11a). This discrepancy is manifested in a monotonously increasing error for the quality of the Ising model fit to the Hill function (Fig.S11b). By comparison, if we plot the ratio of the Ising and Hill fitting errors to the growth-rate data as function of the Hill exponent (Fig.S11c), there is no monotonous increase. This may suggest that the experimental data is indeed not exactly Hill-like.

### S10. BISTABILITY AND CHOICE OF THE SOLUTION BRANCH FOR ISING MODEL FITS WITH $J_0 > J_c$.

In Fig.2 (and Fig.S10) we plot the fitted Ising model interaction parameter $J_0$, and we find few cases where this value is significantly larger than the critical value of $J_c = 4$. When this occurs, there is a large range of antibiotic concentrations for which there are two branches of solution of Eq.(11). We now inspect this bistability in more detail.

In Fig.S12(a) we show this region by plotting the two branches of stable solutions, and the unstable solution, for the growth-rate of *E. coli* as function of the concentration of Ampicillin and Carbenicillin. These antibiotics were fitted with very large values of $J_0 = 7.795, 7.908$, respectively.

In Fig.S12(b) we show the detailed comparison of the fitting of the two branches to the growth-rate data, and the corresponding comparison of the calculated variance (Eq.12) to the measured PGRH. While the fitting to the growth-rate seems to be comparable using either one of the Ising solution branches (although they have distinctly different behavior on either side of the sharp drop), the lower branch (left panels, blue lines) seem to provide a much better fit for the PGRH. Specifically, the observed PGRH seems to have some of the asymmetric behavior predicted by the Ising model, with a low PGRH on the low concentration side of the drop, and much higher on the high concentration side (while its exactly opposite for the upper branch solution).

### S11. COMPARISON OF THE ERRORS OF THE FITS OF THE HILLS AND ISING FUNCTIONS TO THE EXPERIMENTAL DATA OF THE GROWTH-RATE AND THE PGRH.

Comparison of the fitting errors for the Hill and Ising models, per each bacteria species, is given in Fig.S13 (same data that was compiled together in Fig.3).

### S12. CONCENTRATION FLUCTUATIONS THAT ARE PROPORTIONAL TO THE CONCENTRATION, COMPARED TO THE EXPERIMENTAL DATA OF THE PGRH.

We have so far considered that the antibiotic concentration fluctuations grow as the square-root of the concentration (Eq.2). This is motivated by Poisson statistics, which is most relevant at low concentrations. We therefore consider also the possibility that the concentration fluctuations may depend linearly on the concentration, and Eq.2 is modified to be

$$\delta g(c) = A|g(c)'|c \tag{S56}$$

Where $A$ is an arbitrary multiplication factor.

In Fig.S14 we plot the errors in the fits to the PGRH data (for *E. coli*), using either the square-root or the linear relation. Except for one antibiotic (Trimethoprin, compare to Fig.S5), there is no significant difference in the quality of the fits.

*
†

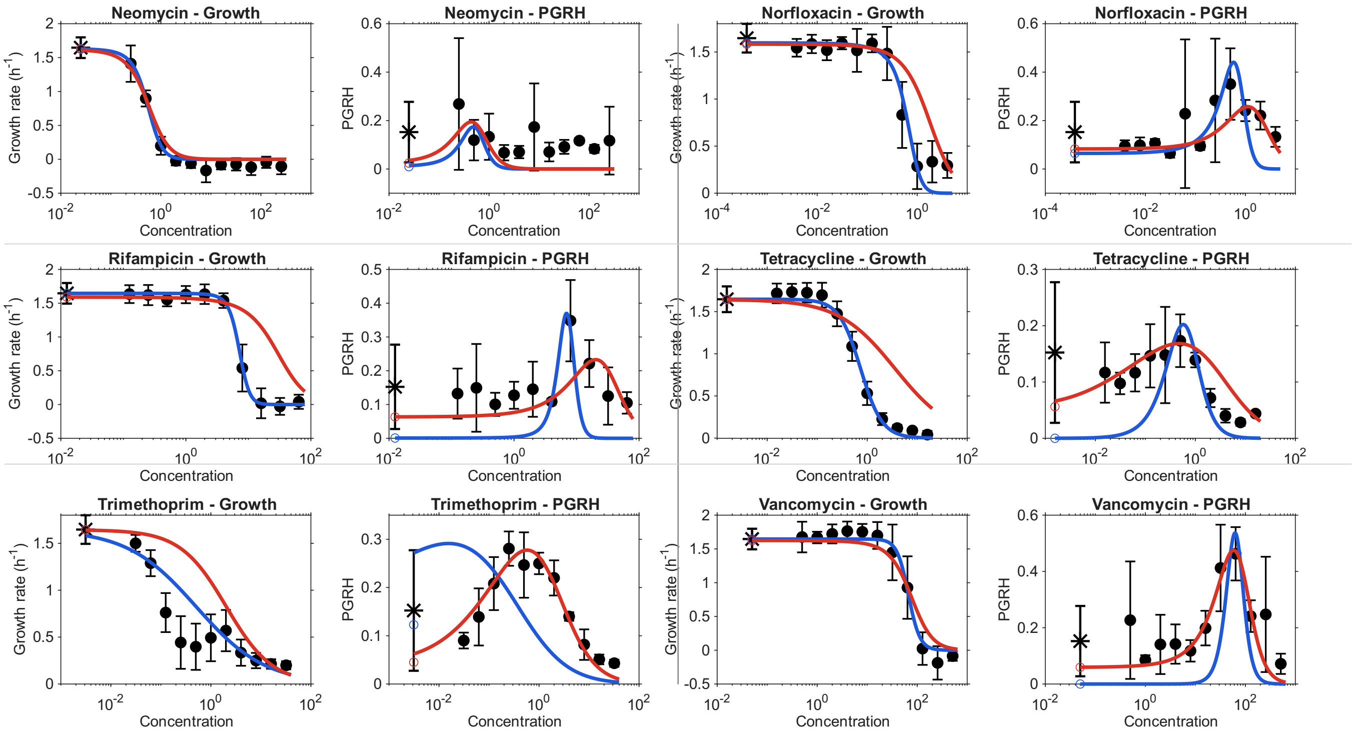


FIG. S2: **Comparison of Hill-model fits to bacterial growth rates and growth-rate heterogeneity (remaining six antibiotics).** The organization and symbols are the same as in Fig. S1. Each antibiotic is represented by two panels: mean growth rate (left) and PGRH (right). Blue curves show fits using only the mean growth-rate data ($W_g = 1$, $W_v = 0$), while red curves show joint fits to growth rate and heterogeneity ($W_g = 1$, $W_v = 1$).

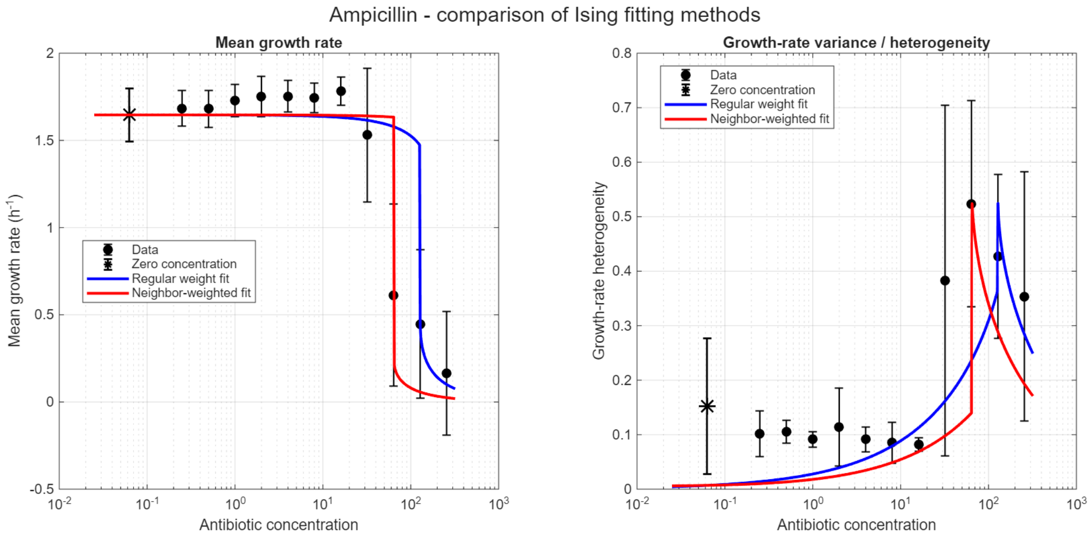


(a)

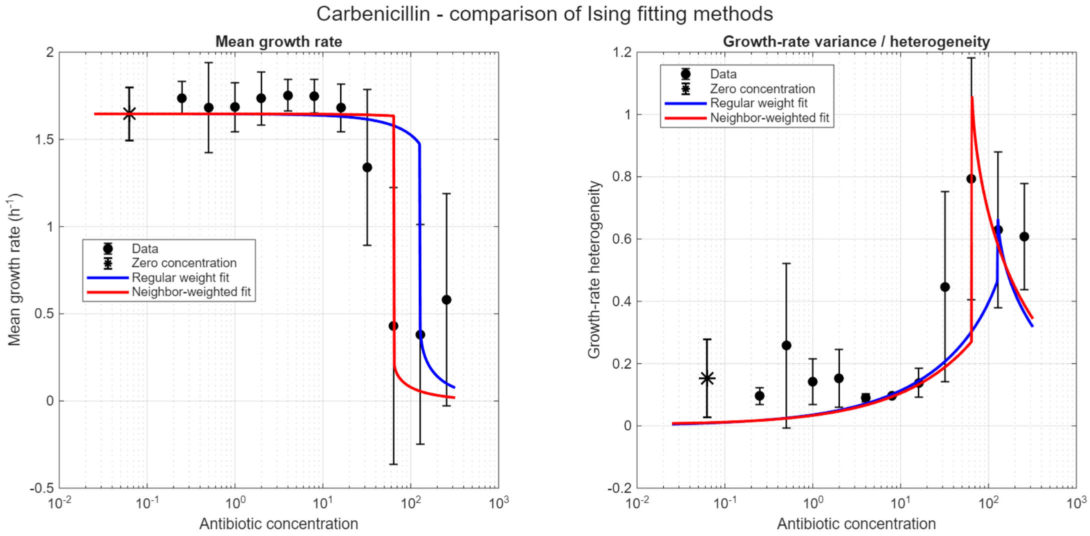


(b)

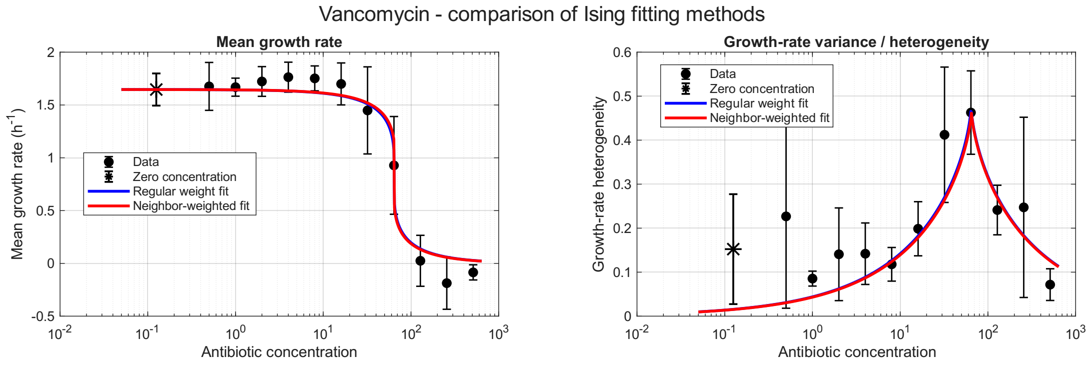


(c)

FIG. S3: Comparison of experimental data (for *E. coli*) and Ising model fits, demonstrating the effect of using in the error calculation weights averaged over three neighboring points of the growth-rate data. The main effect is to move the sharp jump in the Ising model curve to lower antibiotic concentrations, which seems more reasonable for the cases of (a) Ampicillin and (b) Carbenicillin. For all the other cases, as shown for example for (c) Vancomycin, the effect is negligible. The fitting parameters, are shown in Table S1.

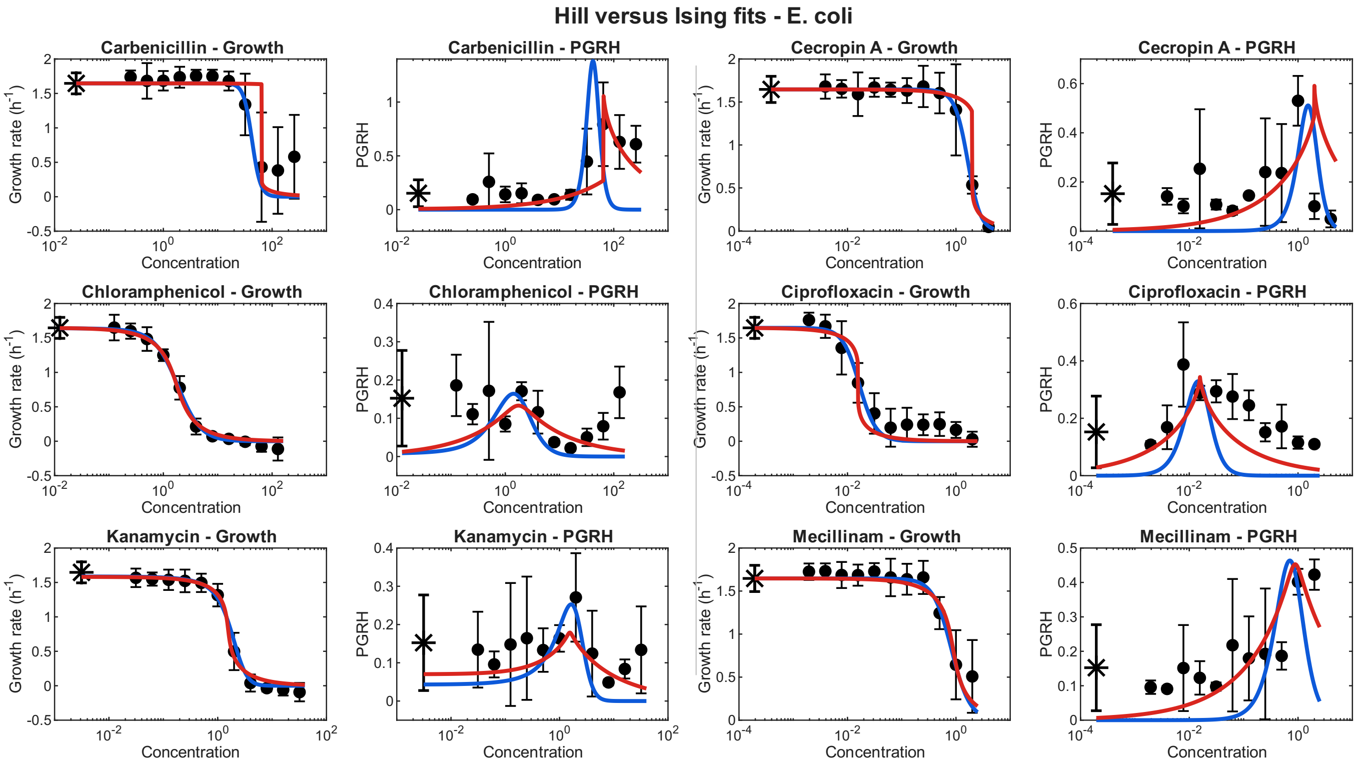


FIG. S4: **Comparison of Hill and Ising models fits to bacterial growth rates and growth-rate heterogeneity, for *E. coli*.** Each antibiotic is represented by two panels: mean growth rate (left) and PGRH (right). Black symbols represent the experimental measurements, with error bars indicating the experimental uncertainties. Blue curves show fits using Hill model, while red curves show the Ising model fits.

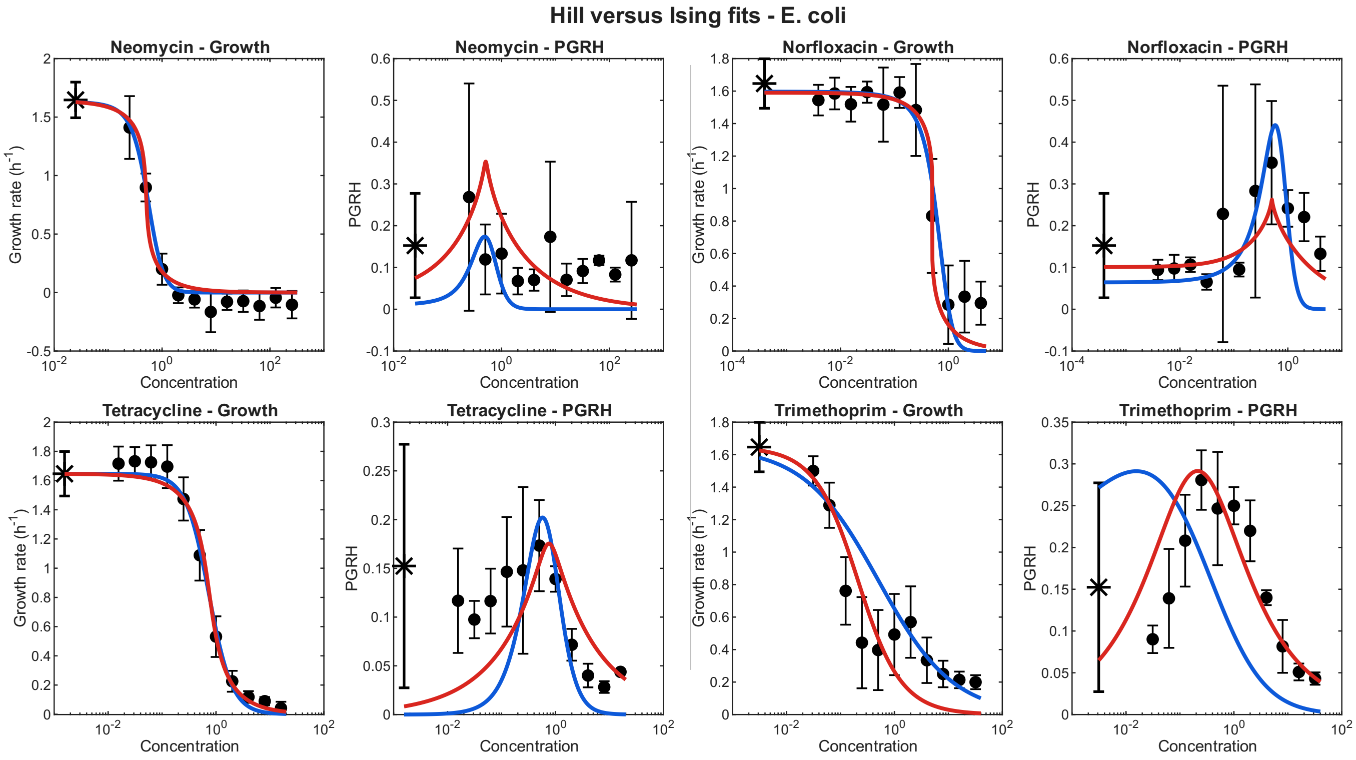


FIG. S5: **Comparison of Hill and Ising models fits to bacterial growth rates and growth-rate heterogeneity, for *E. coli*.** The organization and symbols are the same as in Fig. S4. Each antibiotic is represented by two panels: mean growth rate (left) and PGRH (right). Blue curves show fits using Hill model, while red curves show the Ising model fits.

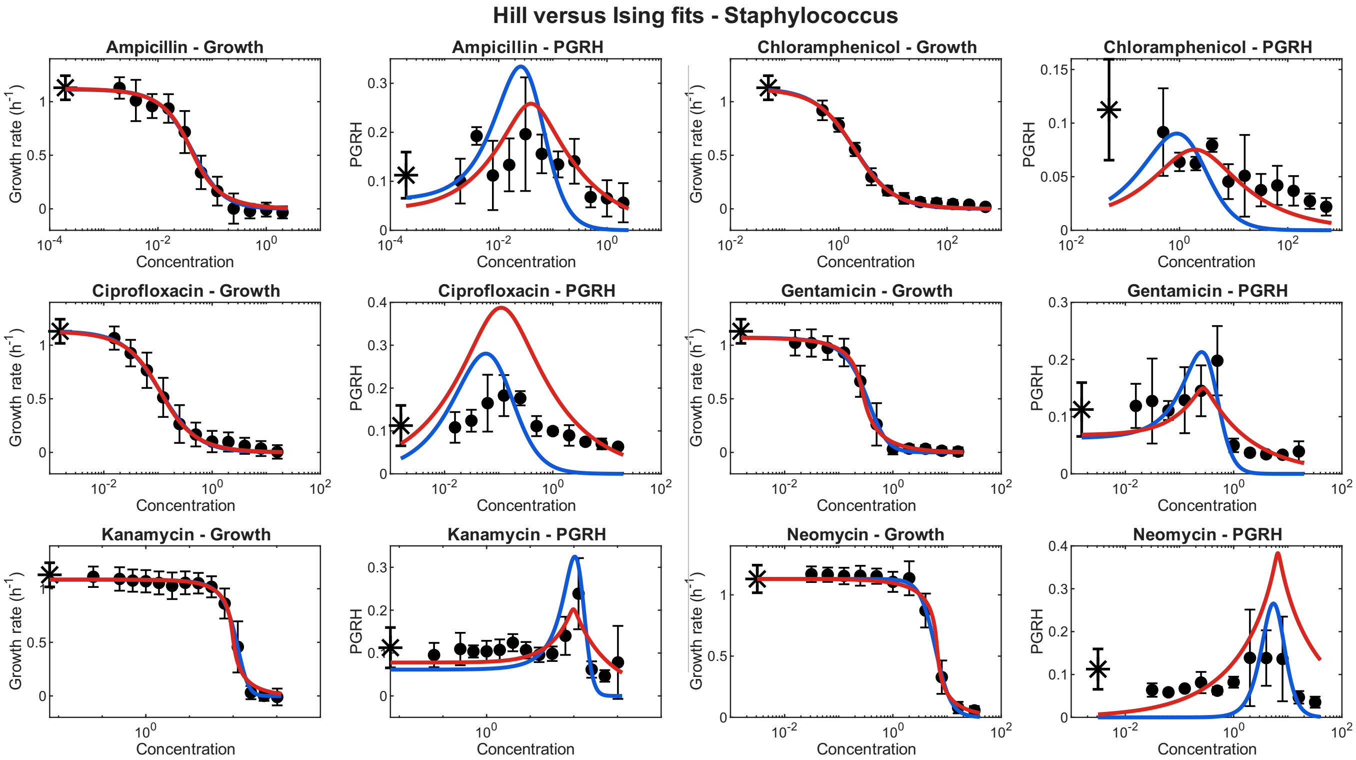


FIG. S6: **Comparison of Hill and Ising models fits to bacterial growth rates and growth-rate heterogeneity, for *Staph.*** Each antibiotic is represented by two panels: mean growth rate (left) and PGRH (right). Black symbols represent the experimental measurements, with error bars indicating the experimental uncertainties. Blue curves show fits using Hill model, while red curves show the Ising model fits.

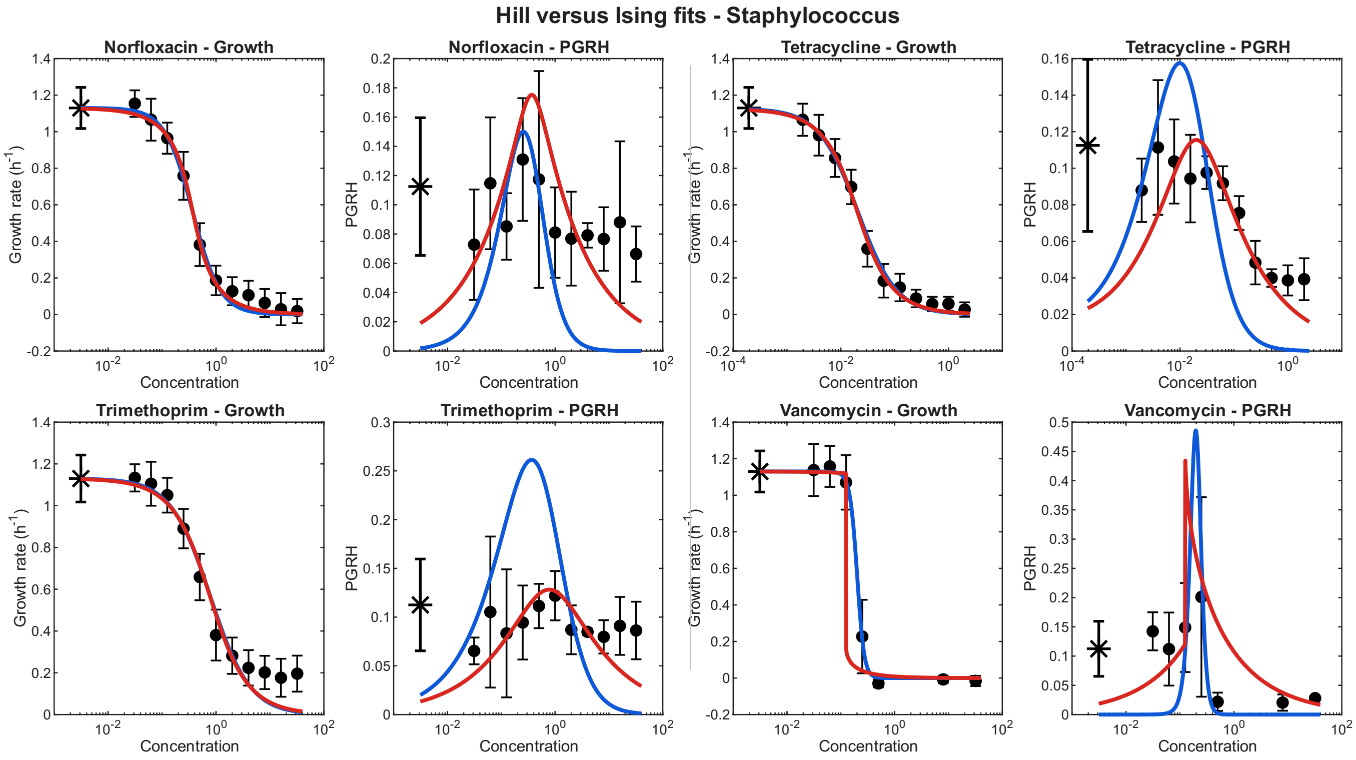


FIG. S7: **Comparison of Hill and Ising models fits to bacterial growth rates and growth-rate heterogeneity, for *Staph.*** Each antibiotic is represented by two panels: mean growth rate (left) and PGRH (right). Black symbols represent the experimental measurements, with error bars indicating the experimental uncertainties. Blue curves show fits using Hill model, while red curves show the Ising model fits.

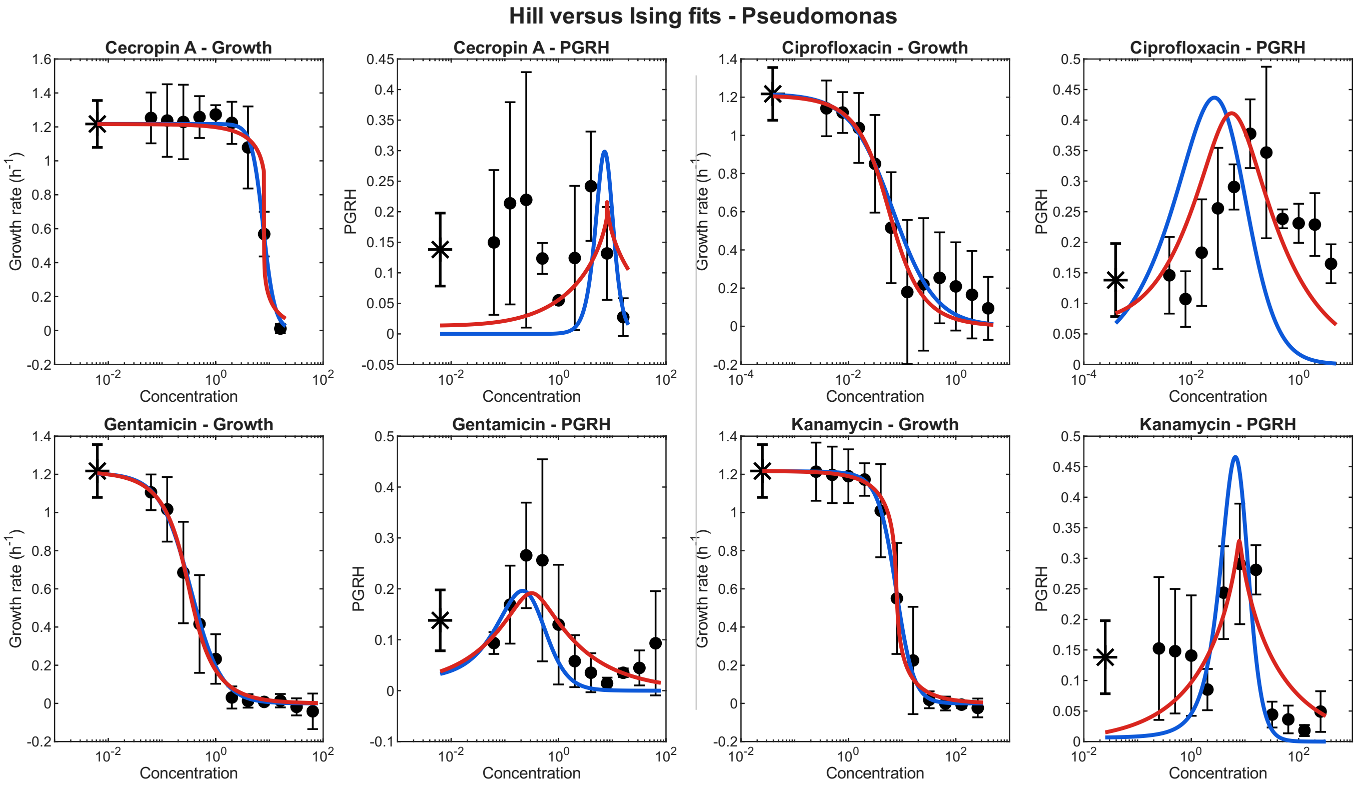


FIG. S8: **Comparison of Hill and Ising models fits to bacterial growth rates and growth-rate heterogeneity, for *Pseu.*** Each antibiotic is represented by two panels: mean growth rate (left) and PGRH (right). Black symbols represent the experimental measurements, with error bars indicating the experimental uncertainties. Blue curves show fits using Hill model, while red curves show the Ising model fits.

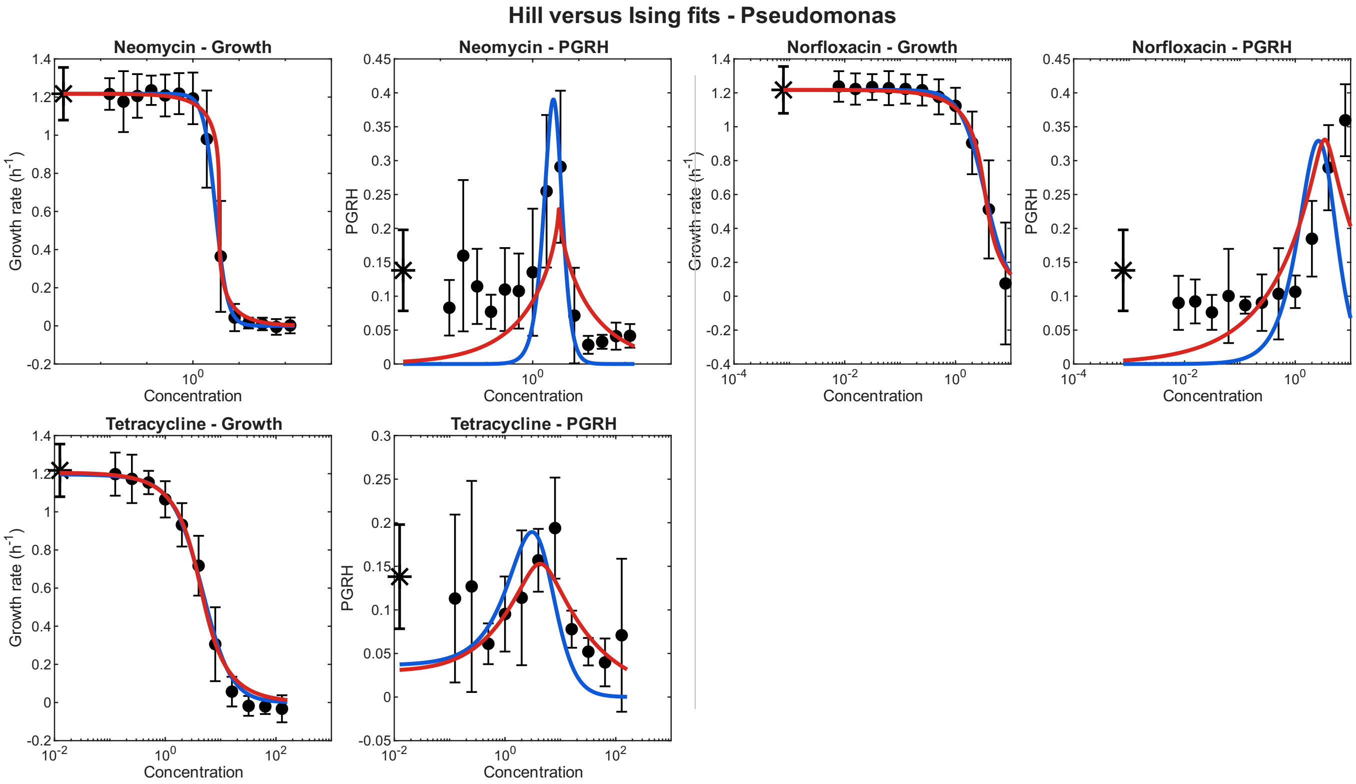


FIG. S9: **Comparison of Hill and Ising models fits to bacterial growth rates and growth-rate heterogeneity, for *Pseu.*** Each antibiotic is represented by two panels: mean growth rate (left) and PGRH (right). Black symbols represent the experimental measurements, with error bars indicating the experimental uncertainties. Blue curves show fits using Hill model, while red curves show the Ising model fits.

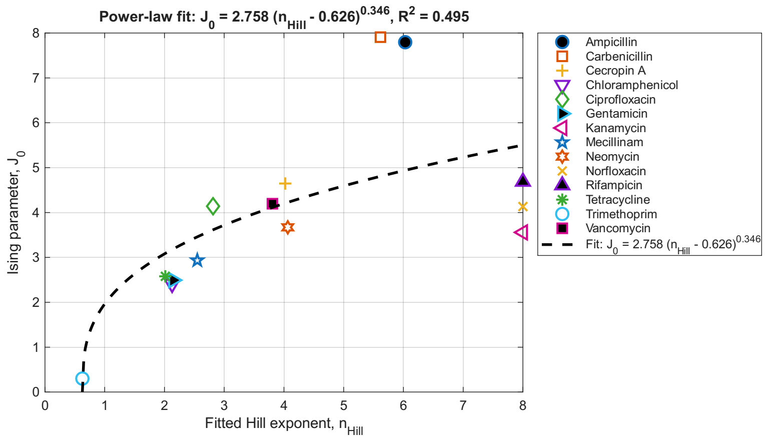


(a)

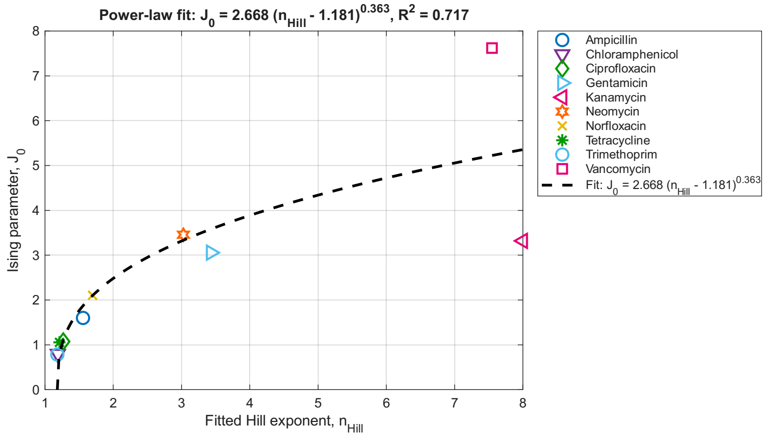


(b)

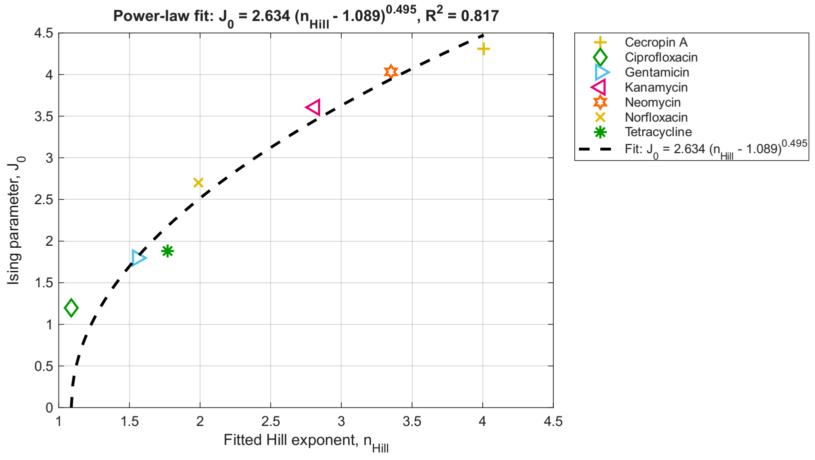


(c)

FIG. S10: Comparison of fitted Hill exponents and Ising interaction strengths. (a) For *E. coli*, the four black-filled symbols correspond to the antibiotics shown in Fig.1. The fits to the other antibiotics are shown in Figs.S4,S5. The dashed line is a polynomial best fit to serve as a guide of the general trend. (b) For *Staph*, and (c) for *Pseudomonas*.

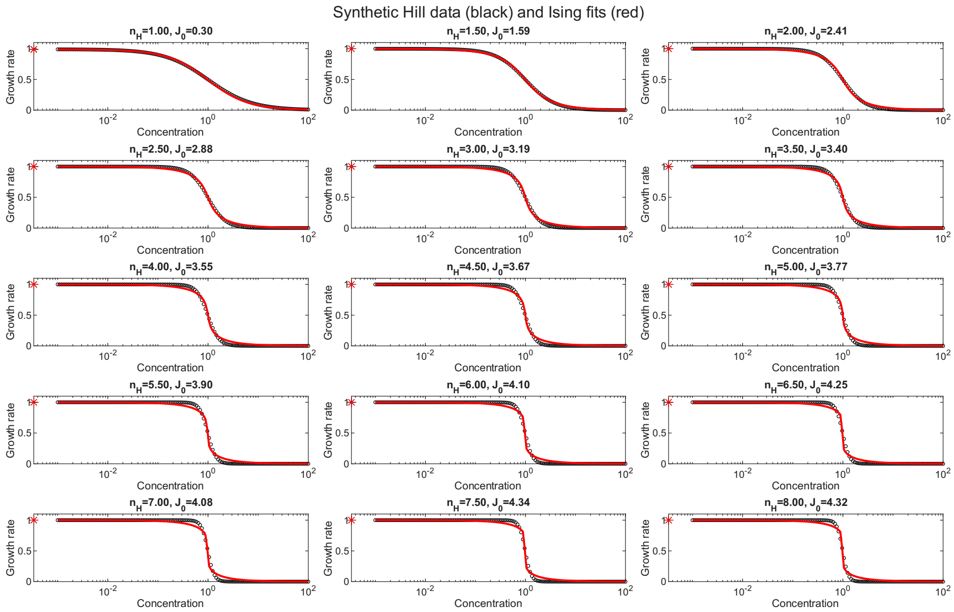


(a)

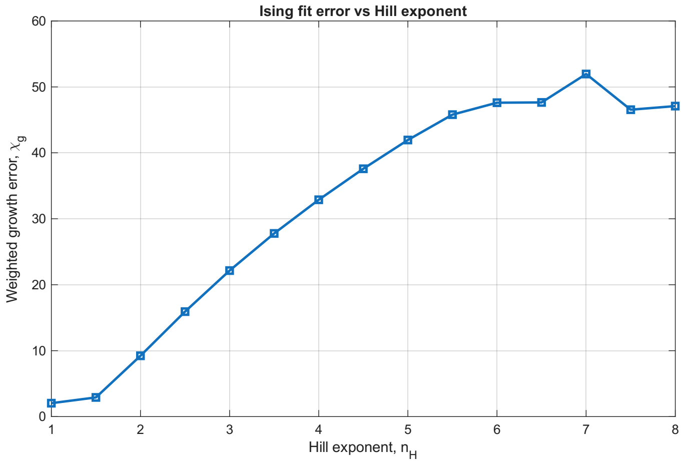


(b)

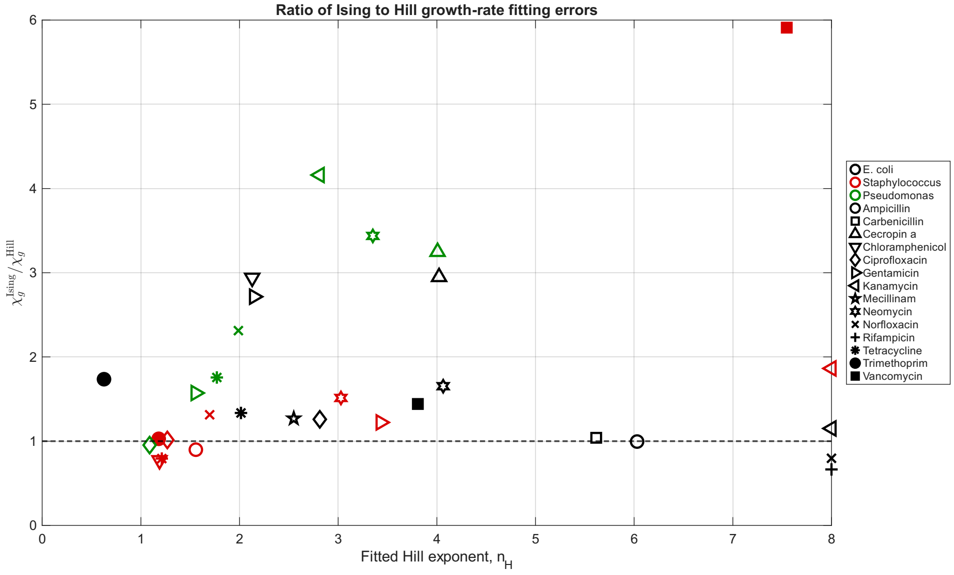


(c)

FIG. S11: (a) Comparison of Hill functions (black circles) to fitting by an Ising model (red line), for different values of the Hill exponent. (b) The error of fitting the Ising model to the Hills function, as function of the Hill exponent. (c) Ratio of the Ising and Hill fitting errors to the growth-rate data as function of the Hill exponent (same data as in Fig.3).

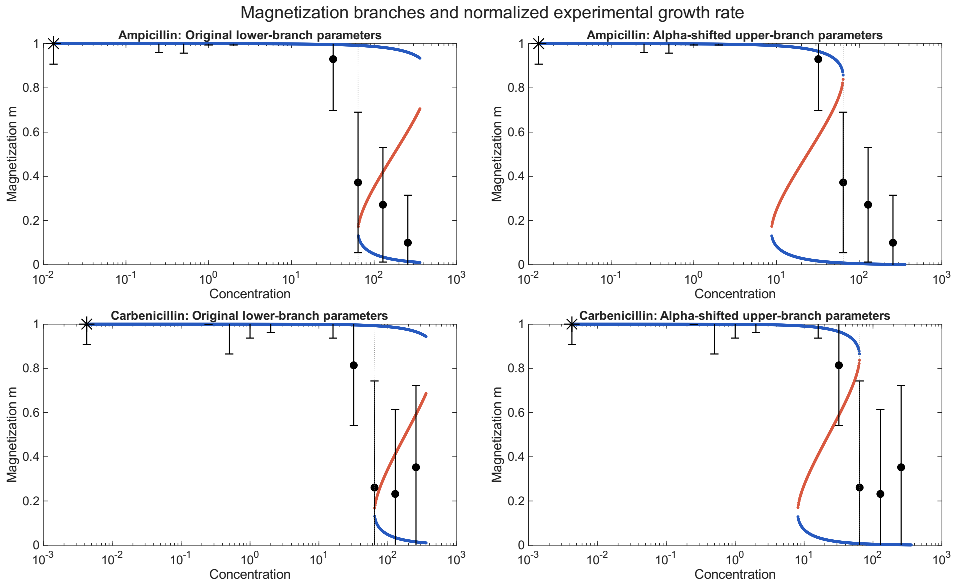


(a)

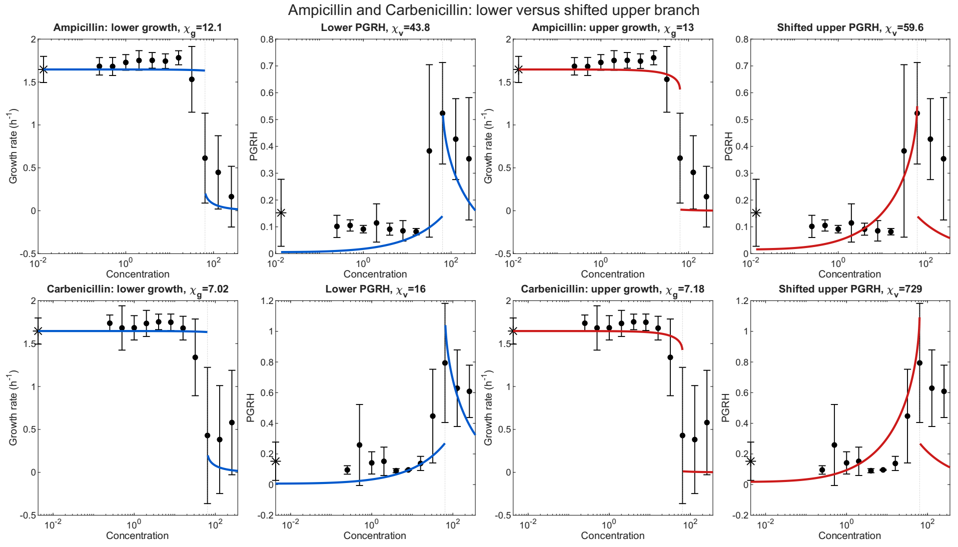


(b)

FIG. S12: (a) Comparison of the growth-rate data of *E. coli* as function of the concentration of Ampicillin and Carbenicillin (black circles and error bars). The lines show the fitted solution of the Ising model, including the bistable region. The left panels show the fit of the sharp drop (vertical dashed line) to the lower branch solution, and the right panels show the fits to the upper branch. Blue lines give the stable solutions and the red line denotes the unstable solution to Eq.(11). (b) Detailed comparison of the fitting of the two branches to the growth-rate data, and the corresponding comparison of the calculated variance (Eq.12) to the measured PGRH.

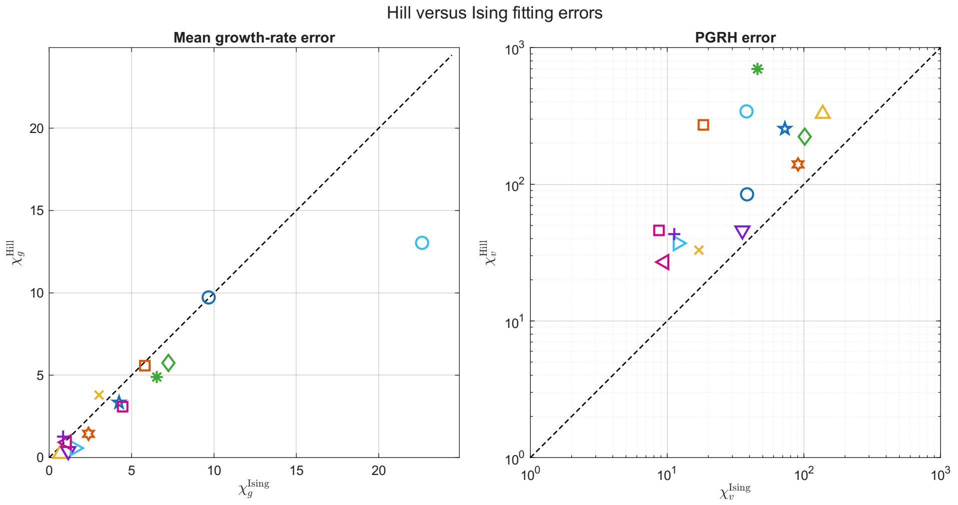


(a)

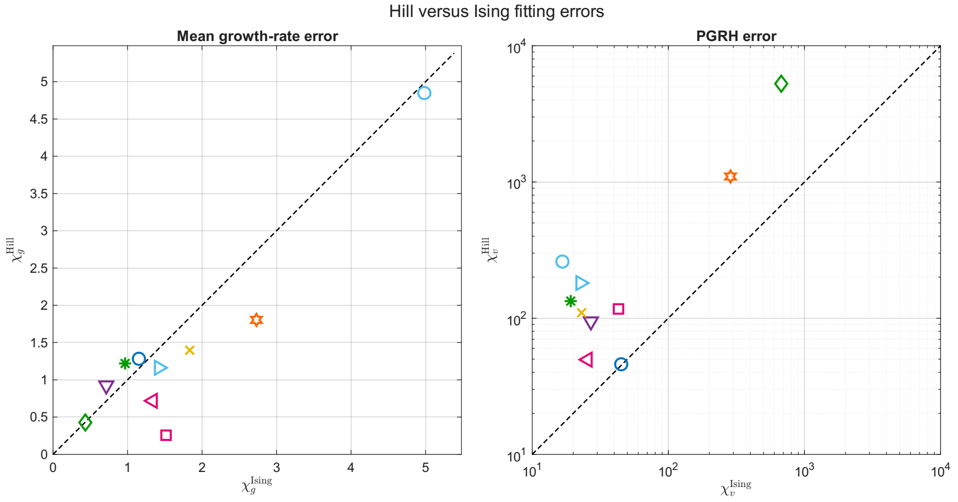


(b)

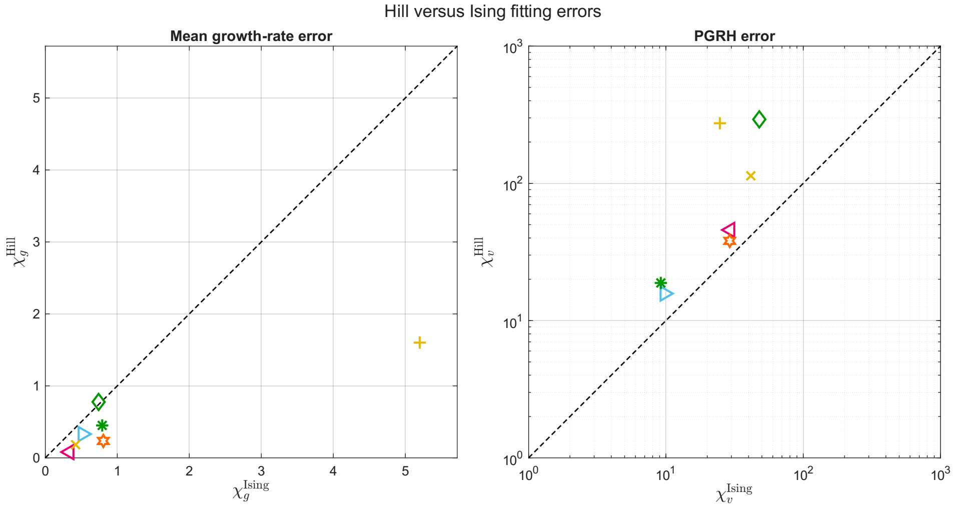


(c)

FIG. S13: (a) Comparison of the fitting errors for the Hill and Ising models for *E. coli*. The dashed line denotes the diagonal where the two errors are equal. Colors and symbols correspond to the different antibiotics as in Fig.S10. (b) For *Staph*, and (c) for *Pseudomonas*.

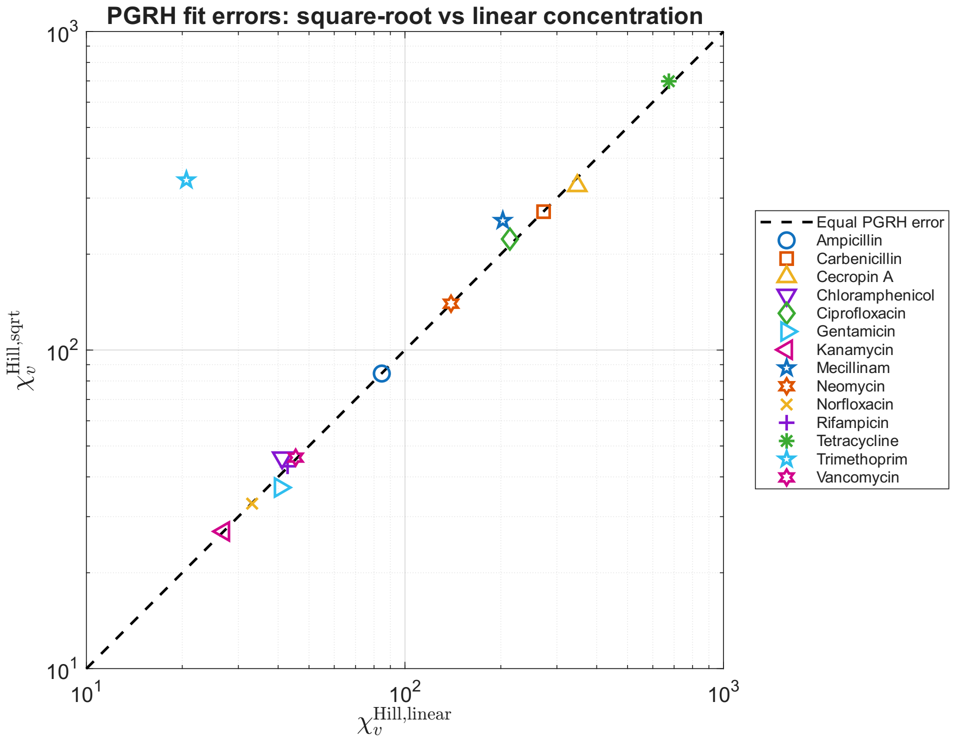


(a)

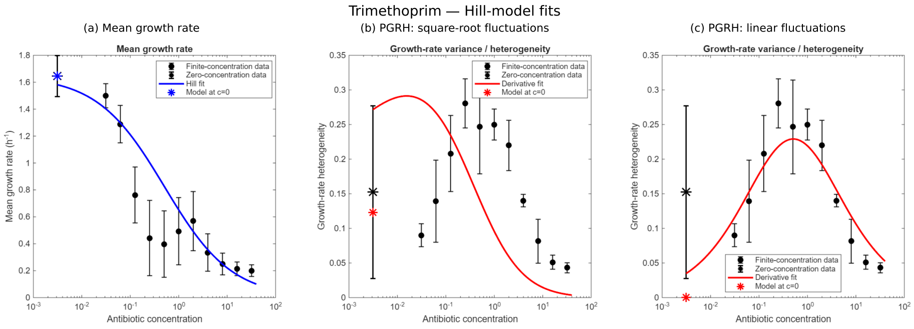


(b)

FIG. S14: (a) Comparison of the fitting errors for the PGRH, for *E. coli*, between square-root and linear concentration fluctuations. The dashed line denotes the diagonal where the two errors are equal. (b) The Hill fit to the growth-rate (left) and to the PGRH (right panels), for square-root (Eq.2) and linear (Eq.S56) concentration fluctuations (compare to Fig.S5).
